# TRIPLE-NEGATIVE BREAST CANCER CELLS RECRUIT NEUTROPHILS BY SECRETING TGF-β AND CXCR2 LIGANDS

**DOI:** 10.1101/2021.01.28.428630

**Authors:** Shuvasree SenGupta, Lauren E. Hein, Yang Xu, Jason Zhang, Jamie Konwerski, Ye Li, Craig Johnson, Dawen Cai, Janet Smith, Carole A. Parent

**Affiliations:** Department of Pharmacology, University of Michigan Medical School; Cancer Biology Graduate Program, University of Michigan; Rogel Cancer Center, University of Michigan; Department of Cell and Developmental Biology, University of Michigan Medical School; Department of Biological Chemistry, University of Michigan Medical School; Life Sciences Institute, University of Michigan

**Keywords:** TNBC, neutrophil, chemotaxis, tumor spheroid, TGF-β, GRO, CXCR2

## Abstract

Tumor associated neutrophils (TANs) are frequently detected in triple-negative breast cancer (TNBC). Recent studies also reveal the importance of neutrophils in promoting tumor progression and metastasis during breast cancer. However, the mechanisms regulating neutrophil trafficking to breast tumors are less clear. We sought to determine whether neutrophil trafficking to breast tumors is determined directly by the malignant potential of cancer cells. We found that tumor conditioned media (TCM) harvested from highly aggressive, metastatic TNBC cells induced a polarized morphology and robust neutrophil migration, while TCM derived from poorly aggressive estrogen receptor positive (ER+) breast cancer cells had no activity. In a three-dimensional (3D) type-I collagen matrix, neutrophils migrated toward TCM from aggressive breast cancer cells with increased velocity and directionality. Moreover, in a neutrophil-tumor spheroid co-culture system, neutrophils migrated with increased directionality towards spheroids generated from TNBC cells compared to ER+ cells. Based on these findings, we next sought to characterize the active factors secreted by TNBC cell lines. We found that TCM-induced neutrophil migration is dependent on tumor-derived chemokines, and screening TCM elution fractions based on their ability to induce polarized neutrophil morphology revealed the molecular weight of the active factors to be around 12 kDa. TCM from TNBC cell lines contained copious amounts of GRO (CXCL1/2/3) chemokines and TGF-β cytokines compared to ER+ cell-derived TCM. TCM activity was inhibited by simultaneously blocking receptors specific to GRO chemokines and TGF-β, while the activity remained intact in the presence of either single receptor inhibitor. Together, our findings establish a direct link between the malignant potential of breast cancer cells and their ability to induce neutrophil migration. Our study also uncovers a novel coordinated function of TGF-β and GRO chemokines responsible for guiding neutrophil trafficking to the breast tumor.

## Introduction

Neutrophils are increasingly recognized as critical tumor-infiltrating immune cells [1]. As part of the innate immune system, they rapidly navigate to sites of infection or tissue injury. Upon arrival to the infected/injured sites, the primary functions of neutrophils are to eliminate invading pathogens and mount an inflammatory response using diverse mechanisms, ranging from phagocytosis and respiratory burst, to the release of toxic granule contents and DNA-containing neutrophil extracellular traps (NET) [2]. Apart from these defense functions, neutrophils have gained attention for their newly discovered role at either promoting [3–5] or hindering [6–8] tumor progression, depending on the types of cancer. The metastasis promoting function of neutrophils during breast cancer have been attributed to the suppression of anti-cancer adaptive immunity [9], stimulation of tumor cell growth [3, 10, 11], or tumor cell invasion and migration ability [4, 12–14]. Tumor-associated neutrophils (TANs) have been detected in varying degrees in the tumor niche of various kinds of malignancies [15], including breast cancer [16, 17]. Yet, cancer-associated guidance cues that drive neutrophil migration to the breast tumor niche are not well defined, partly because of the molecular heterogeneity among different breast cancer subtypes [18].

The major molecular subtypes of breast cancer are luminal A, luminal B, HER2 positive, and triple-negative breast cancer (TNBC), which widely differ in the expression of estrogen hormone receptor (ER), progesterone hormone receptor (PR), and overexpression of human epidermal growth factor receptor-2 (Her-2/neu) [18]. The predominant luminal subtypes are characterized by the presence of hormone receptors (HR+), and exhibit a relatively less aggressive clinical outcome [19]. In contrast, TNBC tumors, which comprise ~15-20% of all breast cancers, are notoriously aggressive with poor clinical outcome, and the occurrence of early distal metastasis is common [20]. Because of the lack of the receptors commonly targeted by existing targeted therapies, the primary therapeutic options for TNBC patients are currently restricted to chemotherapy. However, recent encouraging outcomes with a subset of TNBC subtypes [21] using immune-based strategies have prompted focus on characterizing the immune landscape of the breast tumor niche more comprehensively. Importantly, accumulating evidence has recently highlighted the importance of myeloid cells in regulating the responses of lymphoid populations [22, 23]. Moreover, recent studies have reported the frequent presence of TANs in the tumor niche of TNBC subtypes [16, 17]. In contrast, relatively less aggressive, hormone receptor-positive (HR+) breast tumors have fewer TANs [16], pointing toward an association between the malignant potential of breast cancer cells and the degree of neutrophil recruitment to tumors.

Neutrophils are specially equipped with the ability to directionally migrate toward chemical guidance cues, a process referred to as chemotaxis. They undergo chemotaxis by sensing gradients of chemical cues that bind and activate their cognate receptors, which induce polarized morphology and directed migration [24]. Both tumor and stromal cells are capable of secreting chemokines, cytokines, and growth factors, which in turn facilitate neutrophil mobilization to the tumor niche [25–27]. However, it remains unknown whether tumor cell intrinsic differences in the expression pattern of guidance cues are the sole determinant of the differential presence of TANs in different molecular subtypes of breast cancers.

In the present study, we compared the neutrophil recruitment abilities of tumor-conditioned media (TCM) derived from breast cancer cell lines representing either TNBCs or estrogen receptor positive (ER+) subtypes that constitute the majority of HR+ breast cancer [28, 29]. We followed neutrophil migration in real time in 3 dimensional (3D) collagen matrices towards TCM and determined how tumor-secreted factors modulate neutrophil velocity and directionality. In addition, by establishing a neutrophil-tumor spheroid co-culture system, we observed the ability of breast cancer cells to directly recruit neutrophils. Most importantly, we characterized the nature of the active factors secreted from breast cancer cells using a number of pharmacological, biochemical, and immunological approaches, and identified the factors responsible for neutrophil recruitment by breast tumor.

## Materials and Methods

### Materials

Formyl-methionyl-leucyl-phenylalanine or fMLF (F3506), DMSO, and Pertussis toxin (PT, P7208) were purchased from Sigma-Aldrich. Human recombinant CXCL8 (CHM349), CXCL1 or GROα (CHM329) and GM-CSF (CYT-838) were purchased from ProSpec.

### Cell cultures

We used a panel of 13 different cell lines of breast epithelial origin in the study. Cell lines of different receptor expression status and malignant potential were cultured using medium and serum described in Table 1. All cell lines were maintained at 37°C with 5% CO_2_ except SUM149 cells, which were maintained with 10% CO_2_ [30]. All cell lines tested negative for mycoplasma contamination using the Mycoalert detection kit (Lonza).

**Table 1.** Characteristics and culture conditions of the cell lines used in this study. ER, estrogen receptor; PR, progesteron receptor; HER2; human epidermal growth factor receptor 2. Media conditions: FBS, fetal bovine serum; HS, horse serum.* Media supplemented with 10 μg/ ml insulin (Invitrogen), 10 ng/ml EGF (Peprotech), 0.5 mg/ml hydrocortisone and 100 ng/ml cholera toxin (both from Sigma Aldrich). ** Media supplemented with 5 μg/mL insulin, and 1 μg/mL hydrocortisone (Sigma-Aldrich). MCF10A and MCF10CA1a were obtained from Karmanos Research Institute; MDA 231-TGL, MDA 231-BrM2-831, MDA 231-BoM-1833, MDA 231-LM2-4175 were purchased from Memorial Sloan Kettering Cancer Center; SUM-149 was from Dr. Sofia Merajver, University of Michigan; BT549, Hs578T and BT474 were purchased from ATCC; MCF-7, HCC1428, and T47D were kindly gifted from Dr. James Rae, University of Michigan.

| Cell lines | ER | PR | HER2 | Malignant potential | Culture Media | Serum |
| --- | --- | --- | --- | --- | --- | --- |
| MCF10A (M1) | - | - |  | Benign | *DMEM F12 | 5% HS |
| MCF10CA1a (M4) | - | - |  | Invasive | DMEM F12 | 5% HS |
| MDA-MB-231 | - | - |  | More Aggressive | DMEM | 10% FBS |
| MDA-MB-231 TGL | - | - |  | More Aggressive | DMEM | 10% FBS |
| MDA-MB-231 BrM2-831 | - | - |  | More Aggressive | DMEM | 10% FBS |
| MDA-MB-231 BoM 1833 | - | - |  | More Aggressive | DMEM | 10% FBS |
| MDA-MB-231 LM2-4175 | - | - |  | More Aggressive | DMEM | 10% FBS |
| SUM149 | - | - |  | More Aggressive | **Ham's F12 | 5% FBS |
| BT549 | - | - |  | More Aggressive | RPMI | 10% FBS |
| Hs578T | - | - |  | More Aggressive | DMEM | 10% FBS |
| MCF-7 | - | - |  | Less Aggressive | DMEM | 5% FBS |
| BT474 | + | + | + | Less Aggressive | RPMI | 10% FBS |
| HCC1428 | + | + |  | Less Aggressive | RPMI | 10% FBS |
| T47D | + | + |  | Less Aggressive | RPMI | 10% FBS |

### Isolation of human neutrophils

Blood was obtained from healthy human male and female subjects aged 19–65 years, through the Platelet Pharmacology and Physiology Core at the University of Michigan. The Core maintains a blanket IRB for basic science studies, where HIPAA information is not required. Therefore, while the IRB-approved Core enrolls healthy subjects that conform to the protection of human subject standards, we did not have access to this information. The samples that we received were fully de-identified. Neutrophils were purified using dextran-based sedimentation followed by histopaque-based density gradient centrifugation, adapted from the method described previously [31, 32]. Endotoxin free (< 0.005 EU/mL) cell culture grade water (SH30529, Cytiva) was used to dilute any reagents used for neutrophil isolation from the blood. All tubes used were coated with 0.5% tissue culture grade BSA (A9576, Sigma-Aldrich) diluted in DPBS in order to prevent non-specific cell activation by adhesion to the tube wall. Isolated neutrophils were cleared off residual red blood cells by ACK lysis buffer (10-548E, Lonza) and resuspended in mHBSS buffer (20mM HEPES, pH 7.4, 150mM NaCl, 4 mMKCl, 1.2mM MgCl2, and 10 mg/ml glucose). Viable cells were counted using 0.01% trypan blue (SV30084.01, Cytiva). Cells were more than 99% viable immediately following isolation. In order to address donor-to-donor variability of neutrophil response, cells were routinely tested for minimum basal activity and a robust response to fMLF stimulation before proceeding with daily studies. A dose dependent conversion from basal circular to elongated polarized morphology of fMLF stimulated neutrophils were confirmed by bright-field imaging of two randomly selected fields using 40x objective lens on a Zeiss Axiovert microscope.

### Tumor spheroid generation

Tumor spheroids were generated using the hanging drop method [33]. Briefly, 1000 cancer cells resuspended in full serum media were seeded in 20 μl drops on the inside surface of petri dish lids. Lids were then inverted and placed onto media filled dishes. Spheroids were grown for 4 days in the incubator at 37°C. Day-4 spheroids, visible as aggregates at the bottom of each hanging drop were harvested and used either for imaging or to generate TCM.

### Spheroid imaging

Tumor spheroids were imaged using a 2-photon microscopy. Briefly, day-4 spheroids were fixed with 4% PFA for 1 hour at room temperature. Spheroids were stained with DRAQ5 (Abcam #108410) (1:1000) overnight at 4°C and then mounted onto poly-L-lysine coated coverslips. To reduce scattered light and improve image quality, the spheroids were optically cleared using the ultrafast optical clearing method (FOCM) [34]. Lastly, the coverslips were mounted onto glass slides using ImmuMount (Thermo Scientific #9990402) with small spacers between the slides and the coverslips to prevent the spheroids from being flattened. The spheroids were imaged at 1040 nm using a C-Plan Apochromat 40x/1.3x oil objective on a Zeiss LSM 780. Image stacks were first processed by Histogram Matching with nTracer-AlignMaster [35] to equate brightness differences among slices within the same stack. A median filter with a radius of 0.5px was then applied to suppress the background noise. Center slices of each image stacks were used to representatively demonstrate the spheroids’ structure from each cell type. Movies traversing the image stacks from top to bottom were also generated.

### Harvesting conditioned media from 2D culture

To generate conditioned media (CM or TCM), cell lines were seeded at a density of 0.15×10^6^/ml in 2 ml (6-well tissue culture plate), 16 ml (T75 tissue culture flask), or 32 ml (T150 tissue culture dish) volume of complete medium for 24 hrs, at which point they reached ~ 70% confluence. The culture medium was then removed and cells were gently washed twice with calcium and magnesium free sterile DPBS (Gibco) to remove left over serum containing media. Cells were then incubated with fresh medium supplemented with reduced serum (1%) (Fig.1A) or without serum (Fig.2A) and incubated for an additional 24 or 48 hrs, respectively. The media was then harvested and filtered through 0.22 μm membrane filter to remove dead cell debris. Aliquots were frozen at −30°C until analyzed.

**Figure 1.**
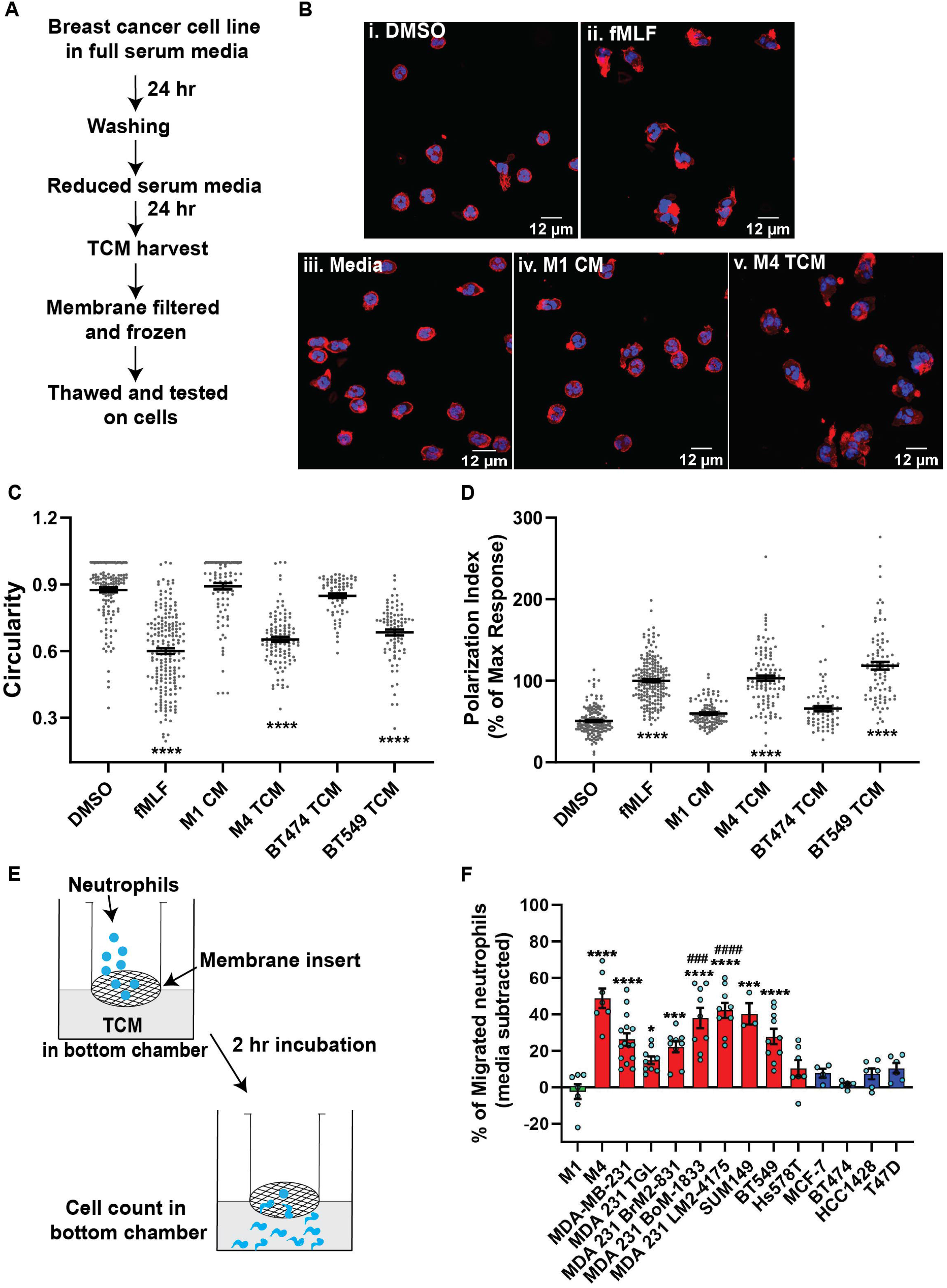
TCM derived from TNBC cell lines induce neutrophil migration. **A**. Flowchart describing how TCM was generated and harvested from cell lines grown under serum reduced conditions. **B.** Representative maximum intensity projection confocal images showing fixed neutrophils stimulated with (i), DMSO, (ii) 100 nM fMLF, equal volume of (iii) media, (iv) M1 CM or (v) M4 TCM, and stained for F-actin with phalloidin-TRITC (red) and nuclei with DAPI (blue). Scale bars are 12 μm. **C&D**. Graphs depicting the circularity (C) and polarization (D) index of neutrophils stimulated with controls or equal volume of TCM derived from different breast cancer cell lines. Each dot represents corresponding value of a single neutrophil for each condition. **E**. Cartoon depicting the neutrophil transwell migration assay. **F**. Graph depicting the percentage of neutrophils that migrated into the bottom chamber of the transwells containing equal volume of CM from M1 cells or TCM from different breast cancer cell lines. Red and blue bars represent values of neutrophil migration obtained with TCM harvested from TNBC and ER+ cancer cell lines, respectively. Data shown are mean ± SEM from N=4 (C, D) or 3-15 (F) independent donors represented by individual dots. Each donor (N) represents an independent experiment. ****P≤0.0001, ***P≤0.001, *P≤0.05 when compared with M1 CM (C, D, F) or #### P≤0.0001 and ### P≤0.001 when compared with MDA 231 TGL (one-way ANOVA with Dunnett’s multiple comparisons test).

**Figure 2.**
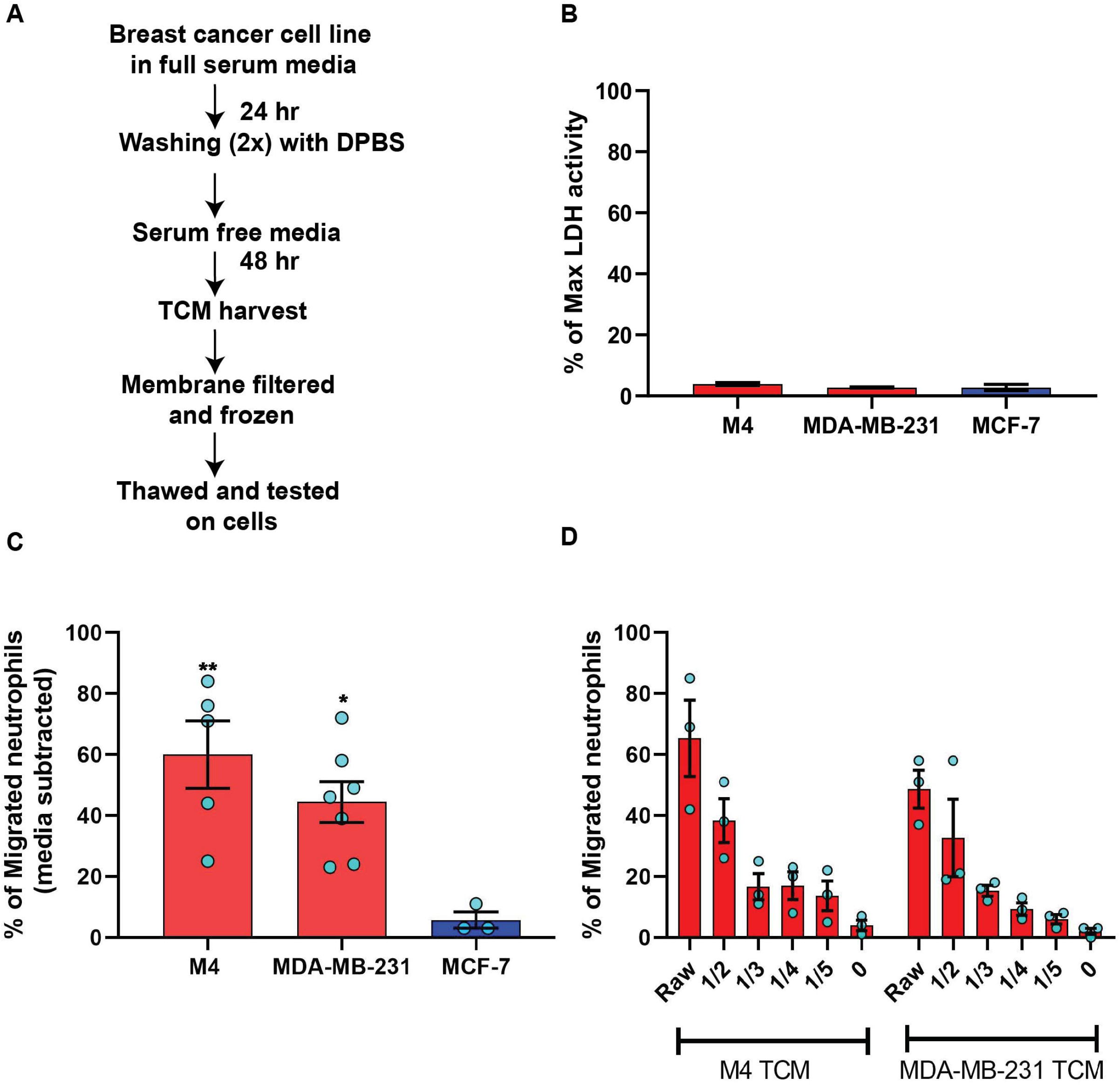
TCM derived from serum deprived TNBC cell lines retain neutrophil recruiting ability. **A**. Flowchart describing how TCM was generated and harvested from cell lines grown under serum free conditions. **B**. Graph depicting spontaneous LDH activity in the supernatant of cancer cell lines grown under serum free conditions measured and normalized with respect to maximum LDH activity. Bars represent mean percentage of max LDH activity ± SEM from 3 independent experiments. **C**. Graph depicting the percentage of neutrophils that migrated into the bottom chamber of the transwells containing equal volume of TCM from different breast cancer cell lines cultured in serum free condition. Red and blue bars represent values of neutrophil migration obtained with TCM harvested from TNBC and ER+ cancer cell lines, respectively. **D**. Graph depicting the percent of neutrophils migrating into the bottom chamber of the transwells containing serially diluted equal volume of TCM from two different TNBC cell lines. Results are presented as mean ± SEM from N=3-7 (C) or N=3 (D) independent donors represented by individual dots. Each donor (N) represents an independent experiment. **P≤0.005, *P≤0.05 when compared with MCF-7 (one-way ANOVA with Dunnett’s multiple comparisons test).

### Harvesting conditioned media from 3D spheroid culture

To generate TCM from cells grown in 3D spheroids, approximately 30 day-4 spheroids were placed in each well of poly-HEMA-coated, 96-well, V-bottom plates. Once settled to the bottom, spheroids were gently rinsed with DPBS and incubated in 200 μl serum-free media for 48 hrs. The media was then harvested and filtered through a 0.22 μm membrane filter. Aliquots were frozen at −30°C until analyzed.

### Neutrophil polarized morphology analysis by immunofluorescence

For the analysis of actin polymerization and polarized morphology, neutrophils were seeded at 1-2×10^6^ cells/ml in 1 ml volume in a 35 mm dish with 20 mm #1.5 coverslip (MatTek). The coverslips were pre-coated with 25 μg/ml fibrinogen (F4883, Sigma-Aldrich) in DPBS for 1 hr at 37°C. Neutrophils were incubated 5-10 min in the incubator at 37°C to allow floating cells to settle down on the coverslip. Cells were then stimulated with an equal volume of TCM, media control, positive control fMLF or vehicle control DMSO and were further incubated for 15-20 min at 37°C. Following stimulation, the entire liquid medium was gently discarded from each dish and the adherent cells were immediately fixed with 4% PFA (Electron Microscopy Sciences), 0.1% glutaraldehyde (G5882, Sigma-Aldrich), 5% sucrose (BP220, Fisher Scientific), and 0.1M cacodylate (11652, Electron Microscopy Sciences), for 15 min at room temperature (RT). Once fixed, cells were washed with 0.1M glycine (G8898, Sigma Aldrich) in DPBS to quench the autofluorescence of glutaraldehyde. Washed cells were next permeabilized with 0.05% saponin (84510, Sigma Aldrich) in 3% BSA-containing DPBS for 15 min at RT and stained with phalloidin-TRITC (P1951, Sigma Aldrich) and DAPI (D9542, Sigma Aldrich). Cells were imaged using a 63x objective lens on a Zeiss LSM880 confocal microscope. Cells in three to five different fields across the coverslip were captured randomly per condition in each experiment. F-actin polymerization and cell circularity were measured using the ImageJ image analysis software. Briefly, integrated density, which is a product of cell area and mean fluorescence intensity, and circularity were obtained from ImageJ after manually identifying the boundary of each cell in a given field. Polarization index was calculated by normalizing the integrated density of each cells with respect to the positive control (fMLF) in each experiment.

### Neutrophil polarized morphology analysis by bright-field microscopy

To analyze cell circularity in response to TCM fractions or TCM in the presence of inhibitors, 0.15×10^6^ neutrophils were seeded in each well of a fibrinogen coated 8 well chambered cover glass with #1.5 cover glass (Cellvis). Cells were allowed to adhere to the cover glass by incubating at 37°C/5% CO_2_ for 5 min followed by stimulation with an equal volume of TCM fractions, controls or TCM with or without the inhibitor for an additional 15 min at 37°C/5% CO_2_. Bright-field images of the cells in two to three randomly selected fields per condition were captured using a 40x objective lens on a Zeiss Colibri microscope. Circularity of each cell was measured by identifying cell boundary manually using ImageJ software.

### Transwell assay

To quantify the ability of TCM to induce neutrophil migration, transwell migration chambers with membrane inserts of 3 μm diameter pore size (Corning and Greiner Bio-One) were used. Both the inserts and the bottom wells were coated with 2% tissue culture grade BSA for 1 hr at 37°C to prevent strong neutrophil adhesion. Coated inserts and wells were rinsed with DPBS twice to remove residual BSA. Freshly isolated neutrophils at a density of 4×10^6^/ml were seeded onto the inserts (100 or 200 μl), and placed in 24-well or 12-well plates, respectively. Control chemoattractants or TCM were gently added in 600 or 1200 μl volume to the bottom wells of 24-well or 12-well plates, respectively, avoiding any bubble formation between the insert and the liquid in the well. Migration was allowed to take in 37°C/5% CO_2_ for 2 hrs. The percentage of neutrophils migrated to the bottom chamber was calculated from the cell counts obtained using a hemocytometer.

### Under agarose assay

Under agarose migration assay was performed following the protocol described elsewhere [36]. Briefly, 0.5% SeaKem ME agarose (Lonza) in 50% each of DPBS and mHBSS was poured and allowed to solidify in 1% BSA-coated 35 mm glass bottom dish with 20 mm micro-well #1.5 cover glass (Cellvis). Using a metallic hole punching tool, three 1 mm diameter wells were carved out at a 2 mm distance from each other. Neutrophils were stained with 0.5 μM CellTracker Red CMPTX dye (Invitrogen) for 15 min at 37°C in rotation and re-suspended in mHBSS. In some experiments, cells were pre-treated with pertussis toxin (PT) for 2 hrs followed by staining in PT supplemented buffer. 7 μl (5×10^4^ cells) of stained cells were added to the wells at the edges and cells were allowed to migrate towards the center well containing 7 μl of either fMLF (100 nM) or 50 μg/ml of GM-CSF for 2 hrs at 37°C/5% CO_2_. End point images of cells migrating towards the chemoattractant were acquired using a 10x objective lens (Zeiss Axiovert microscope).

### 3D collagen matrix assay

Migration in 3D matrix of type I collagen was performed as previously described [37]. Briefly, neutrophils were fluorescently labeled by staining with 0.5 μM CellTracker Red CMTPX dye (Invitrogen) for 15 min at 37°C in rotating condition. Collagen mix was prepared on ice by adding sodium bicarbonate (S8761, Sigma Aldrich), MEM (10X, M3024, Sigma Aldrich), and Purecol^®^ (3 mg/mL, 5005, Advanced Biomatrix) in a 1:2:15 ratio. Labeled cells were washed and resuspended in 1x mHBSS. 0.5×10^6^ labeled cells were gently added to the collagen mix such that the final concentration of collagen is 0.2%. The collagen-cell mix was then immediately added to fill two-thirds of a migration chamber made of coverslip and glass slide. The slide with the chamber was then placed in upright position in the incubator at 37°C/5% CO_2_ for at least 20 min to allow for the gel to polymerize. The upper one-third empty space of the chamber was filled with chemoattractant, either TCM or controls such as media, CXCL8 or buffer. The chamber was immediately sealed with paraffin mix. Time-lapse video was captured for 40 mins at different positions (see Fig. 3A) across the boundary of the chemoattractant and collagen-cell mix using an environmental controlled Zeiss LSM 880 at 10x magnification with the pinhole open. The Manual Tracking plugin in ImageJ software was used to analyze the migration data, where about 70-100 cells were manually tracked over the 40 min time course in each of three randomly selected positions per condition in each experiment. For CXCL8 and buffer control, cells located 1 to 1.5 mm below the matrix boundary were tracked as cells near the boundary mostly exhibited random slow migration, possibly because high concentration of CXCL8 in the boundary saturates the receptors (CXCR1/2). In contrast, cells in the immediate vicinity of the boundary (~500 μm) showed maximum directional movement for TCM and were tracked for both TCM and media control. The software quantified the velocity and directionality of migration, where the directionality was represented by X-FMI, which reflects the efficiency of forward migration of the cells parallel to the gradient [38].

**Figure 3.**
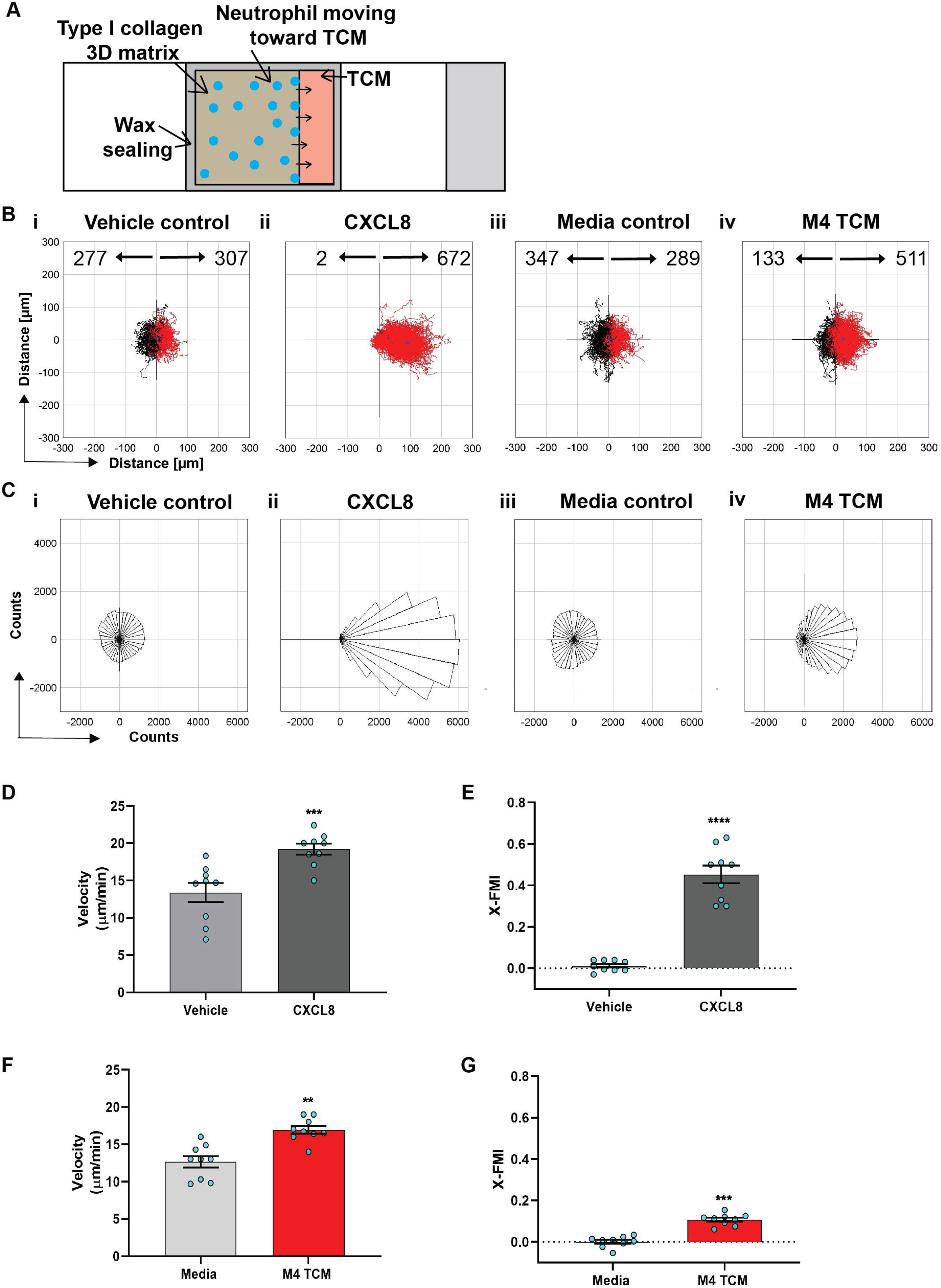
Neutrophils directionally migrate towards secreted factors harvested from TNBC cell line. **A**. Cartoon depicting the migration chamber used to assess the chemotaxis of neutrophils in 3D collagen matrix towards TCM. **B&C**. Plots showing individual cell tracks (B) and rose plots showing angle histograms of the migrating cells (C) generated from the combined tracks from N=3 independent donors in response to vehicle control (Bi, Ci), 500 nM CXCL8 (Bii, Cii), media control (Biii, Ciii), and M4 TCM (Biv, Civ). Numbers with left and right arrowhead in B represent number of neutrophils moving away and towards different conditions, respectively. **D,E,F,&G**. Graphs depicting velocity (D, E) and X-FMI (F, G) of neutrophils migrating towards 500 nM CXCL8 and vehicle control (D,E) or M4 TCM and media (F,G). Each bar represents mean ± SEM from N=3 independent donors with each dot representing average values from independent positions. **** P<0.0001, ***P≤0.0005, **P≤0.005 when compared with corresponding control (paired sample t-test).

### Tumor spheroid-neutrophil co-culture assay

To evaluate neutrophil recruitment toward cancer cells in real time, a co-culture model system was developed (Fig. 5A). Wells of a 24 well glass-bottom plate (Cellvis P24-1.5H-N) were coated with 200 μg/ml fibronectin (Sigma-Aldrich F1141) for at least 1 hr at 37°C. Approximately 20 spheroids resuspended in 250 μl fresh media (Table 1) supplemented with reduced serum (1%) were plated in each well. Spheroids were allowed to attach to the plate for about 4 hrs in the incubator at 37°C/5% CO_2_. Towards the end of incubation, neutrophils were primed with 5 ng/ml GM-CSF for 30 mins. 5×10^4^ or 1×10^5^ primed neutrophils were seeded to each well in 250 μl of 1% serum media. Multiple spheroids in each well were imaged for 24 hrs using an environmental controlled Zeiss Colibri microscope at 10x magnification. To analyze the migration of the neutrophils, the paths of at least 40 neutrophils per video, spaced out both spatially and temporally, were manually tracked using the Manual Tracking plugin in ImageJ software. The position data acquired from the tracking were used in a custom MATLAB script to quantify X-FMI.

### Treatment with pertussis toxin

Neutrophils were treated with pertussis toxin as previously described [39]. Briefly, 4×10^6^ neutrophils/ml were pre-incubated in mHBSS with 10% heat inactivated fetal bovine serum without or with 500 ng/ml pertussis toxin for 2 hours at 37°C with gentle rotation. After 2 hrs, the cells were centrifuged and the pellet was resuspended in mHBSS with 0.5% BSA with or without pertussis toxin. The cells were then loaded into transwell inserts and their migration was assessed as described above.

### TCM fractionation by size exclusion chromatography

30 ml serum free TCM was collected from M4 and MDA-MB-231 cell lines as described above. TCM was concentrated 100-120 fold using Amicon Ultra-15 centrifugal tubes (Millipore Sigma) of 3 kDa molecular weight cut-off. Briefly, 15 ml TCM was centrifuged at 4000 rpm for 40 min at RT. Filtrates were discarded and an additional 15 ml TCM was added to the resulting concentrate, mixed well and centrifuged as before. The final concentrate was buffer exchanged twice with sterile DPBS to get rid of the residual media components and frozen at −30°C until use. Concentrated TCM was applied to a 30 cm x 10 mm Superdex 75 10/300 column (GE Life Sciences) equilibrated with 30 ml DPBS. The column was calibrated using four standards: aprotinin MW 6.5 kDa (Sigma-Aldrich, A3886), cytochrome c MW 12.4 kDa (Sigma-Aldrich, C7150), carbonic anhydrase MW 29 kDa (Sigma-Aldrich, C7025), and bovine serum albumin MW 66 kDa (Sigma-Aldrich, A8531). Sample was run at a rate of 0.2 ml/min and fractions of 150 μl were collected using DPBS as an eluent. Fractions were stored at 4^°^C until use.

### Multiplex ELISA array

TCM concentrates were sent to RayBiotech, Inc. (GA) to screen for a panel of potential neutrophil chemoattracting factors encompassing CXCL chemokine family members CXCL1/2/3 aka GRO, CXCL5, CXCL6, CXCL8, CXCL12, CCL family members CCL2, CCL3, CCL5, growth factors G-CSF, GM-CSF, and cytokine TGF-β. The presence of the analytes was detected and quantified using a high throughput custom Quantibody multiplex enzyme-linked immunosorbent assay (ELISA) (RayBiotech) and the data were analyzed by RayBiotech, Inc. Briefly, concentrates were prepared as described above using 60 ml serum free TCM harvested from the respected cell lines such that the protein concentration approximated 1-2 mg/ml, as estimated using the Pierce BCA protein assay kit (Thermo Fisher). Concentrated samples were tested without dilution in quadruplicate, as per the manufacturer instructions. The protein concentration for each target analyte was quantified by comparing the fluorescence signal from the sample with the individual standard curve generated by known concentrations of each protein.

### Western blotting analysis

ERK phosphorylation was assessed using western blotting analysis. In brief, neutrophils were pretreated with 2 mM pefabloc protease inhibitor (Sigma Aldrich) for 15 min at 37°C with gentle rotation. 4×10^6^ neutrophils/ml were seeded in 500 μl in each well of a 12 well tissue-culture plate pre-coated with 25 ug/ml fibrinogen. Cells were allowed to settle down in the well by incubating the plate for 5 min at 37°C. Cells were then exposed to an equal volume of TCM, media control, positive controls including fMLF and CXCL1 or the corresponding vehicle controls. After 5 min stimulation at 37°C/5% CO_2_, the plate was placed on ice and the cells were lysed in 1X Laemmli buffer (Biorad) containing 5% β-mercaptoethanol (VWR), phoSTOP (Millipore Sigma) and Halt^tm^ protease and phosphatase inhibitor cocktails (Thermo Scientific) at 95°C for 10 min. An equal volume of each was resolved by 10% SDS–PAGE, transferred onto PVDF membranes (Millipore), blocked with 5% nonfat dry milk for 1 hr and probed with primary Abs: phospho-p44/42 MAPK (ERK1/2) (Thr202/Tyr204, dilution 1:2000, Cell Signaling Technology cat. no. 4370) and alpha tubulin (dilution 1:2000, Proteintech Cat no. 66031-1-Ig). Bands were visualized using HRP-conjugated secondary antibodies (dilution 1:8000, Jackson ImmunoResearch), SuperSignal™ West Pico PLUS Chemiluminescent Substrate, and a C600 digital imaging system (Azure). Integrated density for each band was measured using ImageJ software.

### Neutrophil inhibition

Neutrophils were pre-treated with the CXCR2 receptor antagonist AZD5069 (Cayman Chemical), the TGF-β1 receptor ALK5 antagonist SB431542 (Cayman Chemical), or the SMAD3 phosphorylation inhibitor SIS3 (Cayman Chemical) at 1 μM, 1 μM and 3 μM, respectively, or with vehicle control DMSO for 30 min at 37°C with gentle rotation before they were added to the polarization or transwell migration assay.

### Lactate dehydrogenase (LDH) assay

Cancer cell membrane integrity was analyzed for the detection of cell death using the LDH cytotoxicity assay kit (Thermo Scientific) based on the manufacturer instructions. Briefly, cancer cells were seeded at 0.15×10^6^ cells/ml in 100 μl volume in a 96-well tissue culture dish in two sets of triplicate wells. The control set was for maximum LDH release by subjecting cells to undergo complete lysis, and the test set was for spontaneous LDH release. Cells were subjected to incubation and subsequent media change as described for generating serum free TCM. Three days after initial seeding, 10 μl of 10x lysis buffer and 10 μl of sterile water were added to the control and test set, respectively. After incubating the plate for 45 min in the incubator at 37°C/5% CO_2_, 50 μl from each well was transferred to a fresh 96 well plate, mixed with 50 μl LDH reaction mix, and incubated for additional 30 min at RT protected from light. Finally, the reaction was stopped by adding the stop solution and absorbance was measured at 490nm and 680nm by a SpectraMax M5 Multi-Mode Microplate Reader (Molecular Devices). Absorbance was corrected by subtracting background signal of the instrument followed by correcting for the absorbance values of corresponding media. The percentage of maximum LDH release was calculated with respect to corresponding control set.

### Cancer cell migration and invasion assay

Endpoint migration and invasion assays were performed using a transwell system in 24-well plates (Falcon, #353504). FluoroBlock™ filter inserts (Corning, #351152) with 8 μm pore size were used that block light transmission from 400-700 nm and allow fluorescence detection only from the cells that migrated/invaded to the bottom side of the inserts with a bottom-reading fluorescence plate reader. The inserts were uncoated for the migration assays, and coated for the invasion assays with 0.2 mg/ml type I bovine collagen PureCol^®^ of which the pH was neutralized to 7.2 – 7.4 using sodium bicarbonate. Cancer cells that were serum starved for 24 hrs were seeded onto the inserts at 0.5×10^5^ cells/ml in 100 μl of serum free media. Cells were allowed to migrate toward the bottom chamber containing 500 μl of full-serum media as the chemoattractant or serum-free media as the negative control. After a 24 hr incubation at 37°C/5% CO_2_, the inserts were transferred to a fresh 24-well plate with black walls containing 500 μl of 4 μM Calcein AM (Biotium) in mHBSS per well. Cells were incubated for 1 hr at 37°C, and the fluorescence reading of migrated/invaded cells was measured from the bottom at wavelengths of 495/515 nm (Excitation/Emission) by a SpectraMax M5 Multi-Mode Microplate Reader (Molecular Devices).

### Bioinformatics

The transcript expression levels of different CXL chemokine family members in breast cancers were analyzed using datasets identified from the cancer microarray database Oncomine (http://www.oncomine.org). A total of 7 datasets from independent studies were selected from a list of studies obtained by setting the search thresholds as follows: Data type, mRNA; gene rank, top 10%; fold change, 2; P-value, 0.0001. Studies that used Affymetrix platforms other than hgu133plus2 and hgu133a were excluded from further analysis. For each of the studies analyzed, data were downloaded from GEO per instructions of the respective publications. Farmer [40], Minn [41], Wang [42], Ivshina [43] used the Affymetrix hgu133A array while Miyake [44], Schuetz [45], Kao [46] used the Affymetrix hgu133plus2.0 array. For the Schuetz dataset [45], we could not get the raw CEL files and subsequently started from the expression matrix provided with the GEO download. CEL files for all other projects were processed using the Robust Multi-array Average (RMA) technique to obtain normalized expression values [47]. Estrogen receptor annotation was obtained from the data download or from Ivshina [43], Wang [42], Minn [41], Miyake [44], Schuetz [45], and Farmer [40] datasets. Estrogen receptor annotation was not provided for Kao [46]. Instead, using simple histograms, the cutoff for ER status was set by visually selecting a point between the bimodal distribution of ER expression; that cutoff was 10. The Limma package of Bioconductor was used to fit linear models to each dataset comparing ER-samples to ER+ [48]. For Schuetz [45], the model controlled for variation from ductal carcinoma in situ (DCIS) and invasive ductal carcinoma (IDC) status. No other study had a co-variate considered. Data for CXCL chemokines were extracted and the log2 fold change between ER− and ER+ were plotted in a heatmap along with the fold change for the datasets respective estrogen receptor 1 (ESR1) gene (probeset 205225_at). Samples (columns) were sorted by ESR1 expression. From the TCGA data portal, we used the search “Project id” IS “TCGA-BRCA” AND “Workflow Type” IS “HTSEQ-FPKM” AND “Data Category” IS “Transcriptome Profiling” to find 1,222 FPKM (fragments per kilobase of exon model per million reads mapped) data files. The log2 of the FPKM values were plotted for our chemokines of interest and samples were ordered by the FPKM of ESR1.

### Statistical analysis

GraphPad Prism software was used for data plotting and conducting statistical analysis by tests that are described in the respective figure legends along with the size of the samples. Tests used included two-tailed paired t test, unpaired t test, 1-way ANOVA with Dunnett’s multiple comparisons test or 2-way ANOVA with Sidak’s multiple comparisons test.

## Results

### Condition media derived from TNBC cell lines induce neutrophil migration

Numerous studies have suggested that soluble factors, such as CXCL chemokine family members, promote neutrophil infiltration in a variety of tumors. For example, CXCL members CXCL8 (aka IL-8) and CXCL1/2 have been implicated in driving neutrophil recruitment in breast cancer [49–52], and overexpression of these chemokines is associated with breast tumor samples and breast cancer cell lines that have negative estrogen receptor expression status and higher aggressiveness [50, 53–55]. We confirmed these observations by comparing the expression pattern of all known CXCL chemokines across breast cancer subtypes that differ in ER expression status by analyzing transcriptome-based expression data identified from Oncomine and TCGA databases (Fig. S1). Out of 12 CXCL chemokines, we found that the expression of CXCL8, CXCL1, and CXCL2 transcripts was significantly elevated in ER− breast cancer specimens compared to ER+ samples in seven, four and three independent studies, respectively (Fig. S1C). Similarly, TCGA data analysis revealed elevated expression patterns of CXCL8, CXCL1, and CXCL5 in ER− breast cancer specimens (Fig. S1D).

The differential chemokine expression pattern detected among breast cancer subtypes prompted us to investigate the neutrophil recruiting ability of a panel of breast cancer cell lines that differ in ER expression status and malignant potential. TNBCs and ER+ breast cancer cell lines along with non-malignant M1 and metastatic derivative M4 cells from the MCF10A breast cancer progression model [56–59] (Table 1) were cultured *in vitro* under identical seeding cell density. The higher malignant potential of the TNBC cell lines compared to the ER+ breast cancer cell lines was confirmed by assessing their migration and invasion abilities (Fig. S2A&B). We harvested TCM after growing cells in serum reduced media for 24 hrs (Fig. 1A), and tested the ability of the TCM to activate neutrophils. Neutrophils stimulated with fMLF, which is a formylated bacterial tripeptide and a potent neutrophil chemoattractant [32], exhibited an elongated morphology with strong F-actin accumulation in the protrusions, detected using fluorescently tagged phalloidin [32] (Fig. 1Bii; quantified in Fig. 1C&D). In contrast, the majority of neutrophils exposed to DMSO control maintained a circular shape similar to resting neutrophils, and a uniform distribution of cortical actin (Fig. 1Bi). Similar to fMLF, TCM derived from metastatic M4 cells induced strong polarized morphology and F-actin depositions at the poles (Fig. 1Bv; quantified in Fig. 1C&D), while most of the cells stimulated with condition media (CM) collected from M1 cells remained circular (Fig. 1Biv; quantified in Fig. 1C&D). In addition, we found a significant decrease in circularity and increase in F-actin intensity of neutrophils stimulated with TNBC BT549 cell-derived TCM, while TCM derived from ER+ BT474 cells failed to induce such changes (Fig. 1C&D). Next, we assessed the ability of the TCM to induce neutrophil migration using a transwell setup (Fig. 1E). Similar to the polarization response, we found that a significant proportion of neutrophils migrated toward TCM derived from metastatic M4 and TNBC cells, but not from ER+ cells (Fig. 1F). Because each breast cancer cell line has a unique genetic background, we also included in our panel bone-, lung- and brain-specific highly metastatic MDA-MB-231 cells selected from the same parental MDA-MB-231 TGL cells [41, 60, 61]. Interestingly, neutrophil migration towards TCM derived from lung- and bone-specific MDA-MB-231 cells was significantly higher than towards TCM from the relatively less metastatic MDA-MB-231 TGL parental line (Fig. 1F).

To minimize neutrophil activation by the presence of serum in the TCM and to mimic the lack of nutrient availability as encountered by cancer cells in an *in vivo* tumor mass, we harvested TCM from cells cultured in 0% serum for 48 hrs (Fig. 2A) – conditions that lead to minimal cell death (Fig. 2B). We found that serum free TCM derived from M4 and MDA-MB-231 cells induced strong neutrophil migration in transwell assays compared to the TCM harvested from MCF-7 cells cultured under the same conditions (Fig. 2C). Notably, a greater percentage of neutrophils migrated toward serum free TCM (Fig. 2C) than to serum reduced TCM (Fig. 1F) derived from either M4 (57% vs. 49%) or MDA-MB-231 (42% vs. 26%) cells. Importantly, a gradual decrease in neutrophil recruitment with serially diluted serum free TCM showed that TCM triggers a dose-dependent response (Fig. 2D). Together, these findings establish that TCM derived from breast cancer cells with triple negative receptor expression status and aggressive malignant potential induce neutrophil polarization and robust migration by secreting specific chemotactic factors. For subsequent studies, we used TCM generated under serum starved conditions from all cell lines.

### Neutrophils directionally migrate towards secreted factors harvested from TNBC cell line

We next assessed the ability of TCM to induce neutrophil migration in a more physiologically relevant interstitial tissue-like microenvironment by monitoring neutrophil motility through 3D matrices of type I collagen and time-lapse imaging (Fig. 3A). By tracking the movement of neutrophils, we generated trace plots (Fig. 3B) and rose plots (Fig. 3C) that depict the behavior of cells in response to M4 TCM or media control. CXCL8 was used as a positive control. We found that neutrophils migrated directionally towards M4 TCM compared to media control (Fig. 3Biii, 3Biv, 3Ciii, 3Civ; see also Movie S1&S2). We also found that neutrophils migrated towards M4 TCM with a significantly elevated velocity and directionality, compared to media control (Fig. 3F&3G). These increases in cell velocity and directionality also occurred in response to CXCL8 (Fig. 3Bi, 3Bii, 3Ci, 3Cii, 3D&3E; see also Movie S3&S4). However, a lower directionality was measured in response to M4 TCM (0.1) compared to CXCL8 (0.45), which could reflect the complexity of TCM composition. Indeed, unlike a single chemoattractant like CXCL8, the TCM includes multiple active factors that can potentially modulate their effect and have different diffusibility through the collagen. Taken together, these results indicate that active factors secreted from TNBC cells induce directional migration of neutrophils.

### TNBC spheroids secrete factors that recruit neutrophils

We investigated the neutrophil recruitment ability of 3D tumor spheroids generated with breast cancer cell lines. First, we generated tumor spheroids with diameters ranging from 200 to 350 μm using the hanging drop method. Using 2 photon microscopy, we assessed the morphology of the spheroids, which differ in appearance, ranging from irregular, loose aggregates (MDA-MB-231 and BT549) to tight, symmetrical spheres (BT474 and M4) respectively (Fig. 4A, Movie S5-S8). We also determined that under our growth conditions, the spheroids did not harbor a hollow or necrotic core at their center (Fig. 4A), ruling out the possible release of damage-associated molecular patterns (DAMPs) from damaged or dying cells at the spheroid core on neutrophil recruitment [62] (Fig. 4A; Movie S5-S8).

**Figure 4.**
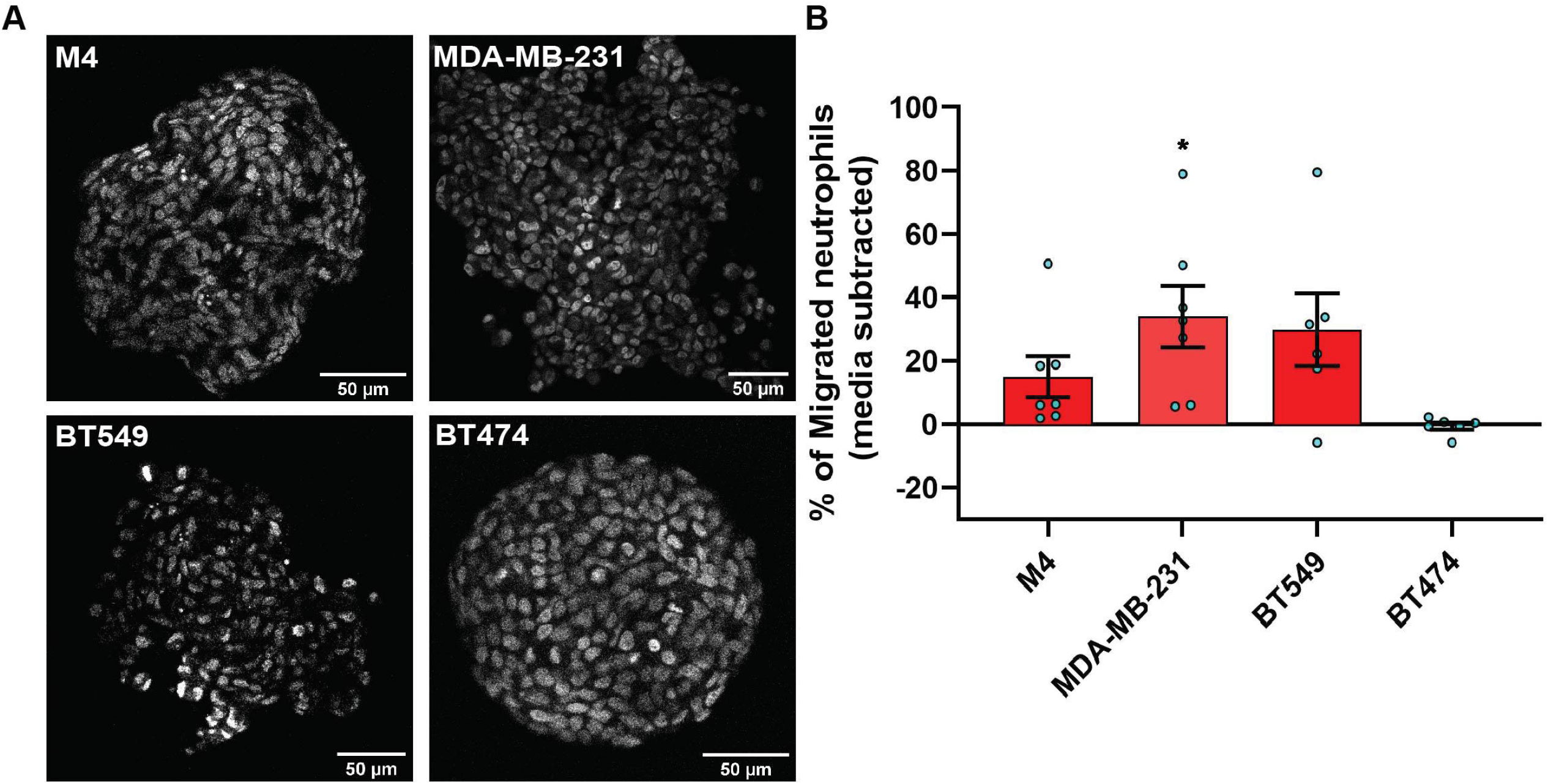
TNBC spheroids secrete factors that induce neutrophil migration. **A.** Representative 2-photon images of day-4 spheroids stained with the nuclear dye DRAQ5. Each image is from the field of view located at the midpoint between the top and bottom of the z-stack imaging range. **B.** Graph depicting the percentage of neutrophils that migrated into the bottom chamber of the transwells containing equal volume of TCM derived from M4, MDA-MB-231, BT549, or BT474 spheroids. Data shown are mean ± SEM from N=6-7 independent donors represented by individual dots. Each donor (N) represents an independent experiment. *P≤0.05 when compared with BT474 TCM (one-way ANOVA with Dunnett’s multiple comparisons test).

To evaluate whether TNBC spheroids secrete neutrophil recruiting factors similarly to TNBC cells grown in monolayers, we assessed neutrophil migration toward spheroid-derived TCM, and found that spheroid-derived TCM from all three TNBC cell lines (M4, MDA-MB-231, and BT549) had neutrophil recruiting activity, while spheroid-derived TCM from the ER+ cell line (BT474) was inactive (Fig. 4B). These results establish that TNBC cells secrete neutrophil-recruiting factors in either 2D or 3D environments.

Next, we developed a neutrophil-tumor spheroid co-culture system to monitor the migration of neutrophils towards tumor spheroids by time-lapse imaging (Fig. 5A). During the early hours of imaging, when neutrophils are not activated and therefore not attached to the surface, we noted that the neutrophils followed the fluid flow and some of them non-specifically got trapped on the surface of the tumor spheroids (Movie S9, S10). However, over time, activated neutrophils clearly attached to the surface, actively migrated towards the M4 spheroids, and accumulated around the spheroid boundary (Fig. 5B, Move S9). In contrast, we found that neutrophils never strongly adhered to the surface or directionally migrated towards BT474 spheroids. Instead, they followed the direction of the flow in the system (Fig. 5C, Movie S10). Indeed, quantification of the directionality of neutrophil migration towards the spheroid revealed a higher directionality towards M4 spheroids, compared to BT474 spheroids (Fig. 5D).Taken together, these findings further affirm that TNBC cells induce directional migration of neutrophils by actively secreting specific chemotactic factors.

**Figure 5.**
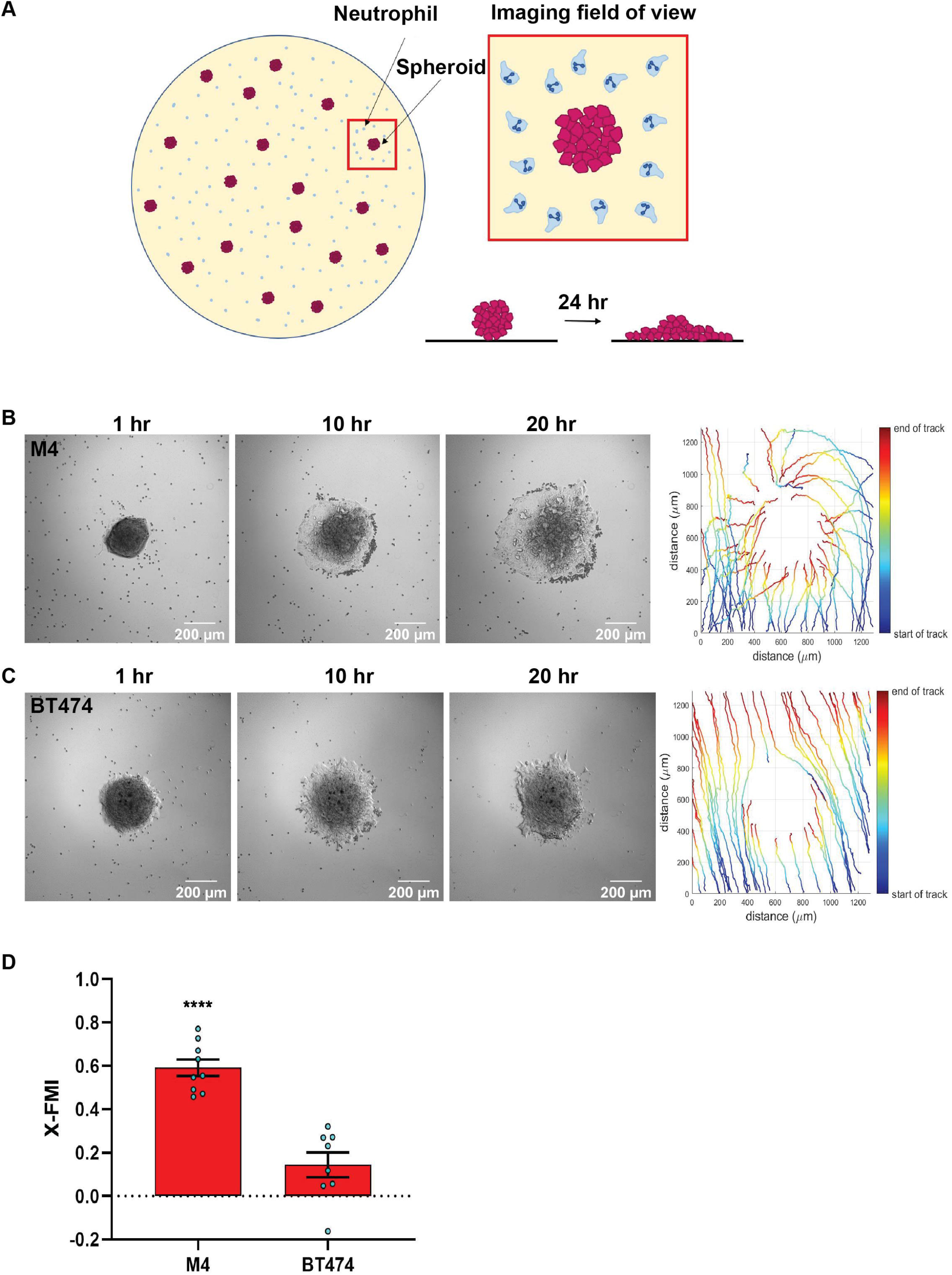
TNBC spheroids actively recruit neutrophils. **A.** Cartoon depicting the co-culture migration system used to assess the chemotaxis of neutrophils towards tumor spheroids in fibronectin coated glass-bottom plate. **B&C.** Representative images of a neutrophil-spheroid co-culture migration assay at 1, 10, and 20 hrs with M4 spheroid (B) or BT474 spheroid (C) at the center of the field of view. Small, dark cells are neutrophils from the same donor. Representative tracks of neutrophils are on the right. **D.** Graph depicting the X-FMI of neutrophils migrating towards M4 or BT474 spheroids. Each bar represents mean ± SEM from N= 3 (BT474 spheroid) or 4 (M4 spheroid) independent donors with each dot representing the average X-FMI for all neutrophils tracked per spheroid. An average of 45 neutrophils were tracked per spheroid. ****P<0.0001 when compared with BT474 (unpaired t-test).

### TNBC cell lines secrete neutrophil activating factors more abundantly than ER+ cancer cell lines

One of the major signaling pathways triggered in neutrophils by a wide range of activating factors, including chemoattractants, is the extracellular signal–regulated kinase (ERK) pathway [63, 64]. ERK pathway is also important for activating the migration machinery of neutrophils [65]. We therefore compared the ability of the secreted factors from different breast cancer cells lines to activate ERK by monitoring its phosphorylation status. We found that TCM derived from either M4 or MDA-MB-231 cells induced ERK phosphorylation, with M4 showing a more dramatic response (fMLF and CXCL1 were used as positive controls) (Fig. S3 and 6A). Importantly, we measured no stimulation with TCM derived from the ER+ MCF-7 cells (Fig. S3 and 6A).

To identify the nature of the activating factors driving neutrophil migration, we first used multiplex ELISA to quantify the relative presence of potential neutrophil chemoattractants /activators associated with cancer, including a number of CXCL and CCL chemokines, cytokines, and growth factors [1, 66] in TCM derived from M4, MDA-MB-231, and ER+ MCF-7 cell lines (Fig. 6B). Among the CXCL chemokines associated with breast cancer of ER− status (GRO, CXCL5, CXCL6, and CXCL8) and breast cancer metastasis (GRO, CXCL5, CXCL6, CXCL8, and CXCL12) [67, 68], we found a greater than 500-fold increase in GRO chemokines (CXCL1, CXCL2 and/or CXCL3) present in the TCM harvested from TNBC cells, compared to TCM harvested from ER+ cells. This high amount of GRO secretion by TNBC cells is in agreement with the robust expression of CXCL1 mRNA in ER− breast tumor specimen compared to ER+ ones ([55] and Fig. S1). We detected a greater amount of CXCL8 present in the TCM from TNBC cells compared to the TCM from ER+ cells, which is in accordance with previous reports [53, 54]. However, compared with GRO chemokines, the fold change in CXCL8 expression was more modest, suggesting that CXCL8 is not the key factor involved in neutrophil activation. In addition, we detected very little to no changes for CXCL5 and CXCL6 in any of the TCM. We also measured > 9-fold increase in CXCL12 in the TCM derived from M4 and MDA-MB-231 cells, compared to MCF-7 TCM. In contrast, the CCL family members were in general less abundantly secreted by breast cancer cells. We detected a 5-fold greater abundance of CCL2 in TCM from TNBC cells than ER+ cells, while very little to none CCL3 in any TCM. In contrast, CCL5 was secreted in equivalent amounts by all breast cancer cells, irrespective of the receptor expression status or malignant potential. Other than chemokines, we detected G-CSF and GM-CSF more abundantly in TCM harvested from TNBC cells compared to ER+ cells. Finally, we found that TGF-β1 was 12-30 fold more abundant in TCM harvested from TNBC compared to ER+ cells. Taken together, these data show that compared to ER+ breast cancer cells, TNBC cells overexpress a number of potential neutrophil recruiting factors, the most abundant of which being the CXCR2 specific GRO chemokines and TGF-β1.

**Figure 6.**
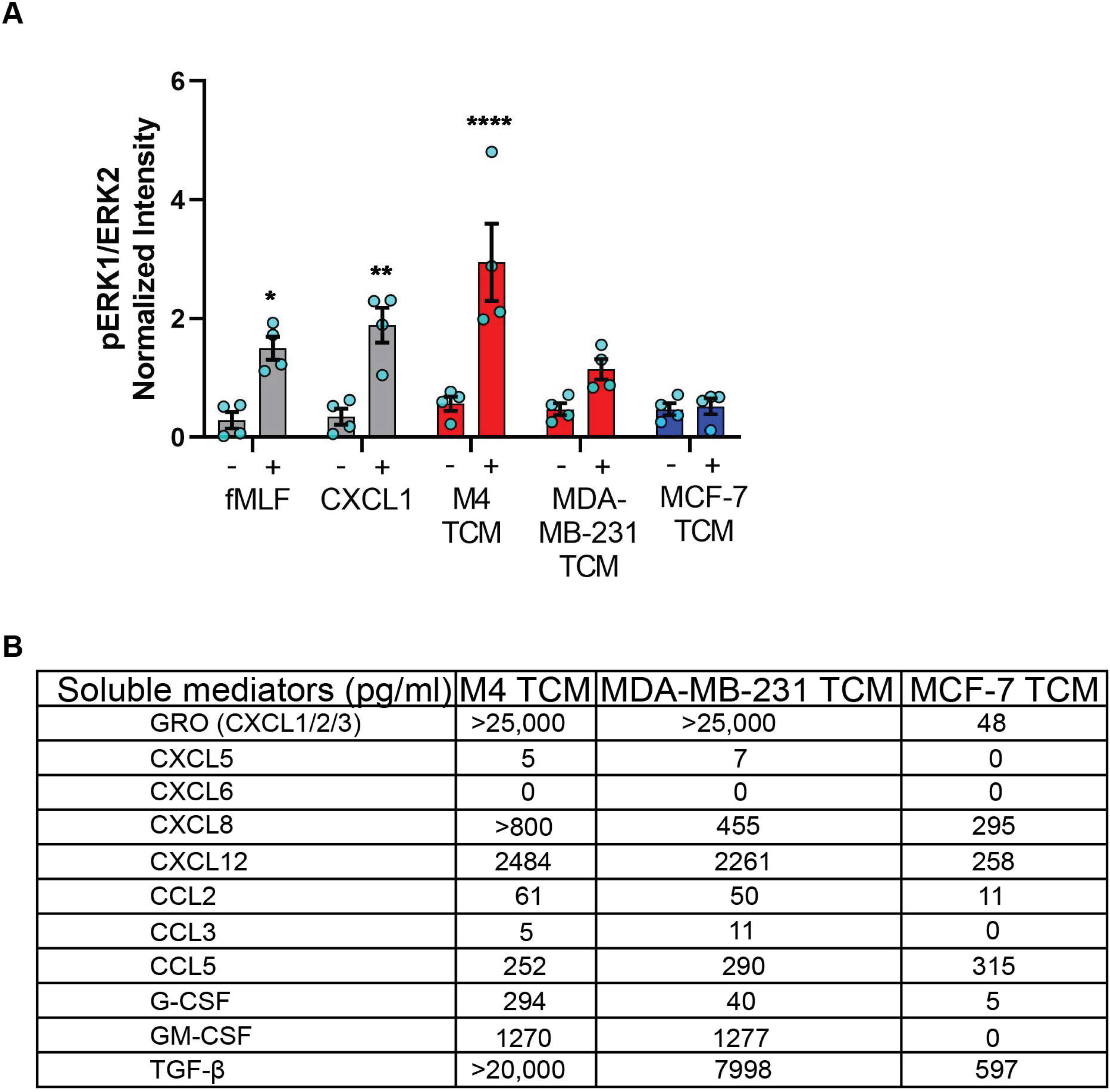
TNBC cell lines secrete neutrophil activating factors more abundantly than ER+ cancer cell lines. **A.** Graph depicting pERK1/ERK2 intensity in neutrophils stimulated with controls (fMLF, CXCL1, media) or equal volume of TCM derived from M4, MDA-MB-231, or MCF-7 cells. The quantified pERK1/2 band intensity was normalized to α-tubulin and is shown as mean ± SEM from N=4 individual donors represented by each dot. **** P<0.0001, **P≤0.01, *P≤0.05 when compared with corresponding vehicle controls (two-way ANOVA with Sidak multiple comparisons test). **B.** Table showing the amount (pg/ml) of chemokines, growth factors and cytokines secreted by M4, MDA-MB-231 and MCF-7 cells determined by multiplex ELISA assay.

### Tumor derived active factors are of ~12 kDa molecular size

To narrow down our search for tumor-secreted active factors, we used a combination of biochemical and pharmacological approaches. We set out to fractionate the TCM based on the molecular weight of their components and identify the fraction with optimum activity. We subjected the concentrates (>3 kDa) of MDA-MB-231 TCM to size exclusion chromatography (Fig. 7A) and tested the resulting fractions for their ability to induce polarized morphology in neutrophils. We found that the fractions carrying proteins corresponding to ~12 kDa molecular weight (fraction “g” in Fig. 7A) induced maximum polarization as indicated by the elongated shape of the cells (Fig. 7Biv). In contrast, most of the cells retained a circular shape when stimulated with fractions containing proteins that were either larger (Fig. 7Bii&iii) or smaller than ~12 kDa (Fig. 7Bv&vi). Indeed, when we quantified the extent of polarization induced by a series of fractions, we found that circularity was significantly reduced when exposed to MDA-MB-231 TCM itself or TCM-derived fractions corresponding to 12 kDa sized proteins (Fig. 7C, fractions e,f,g,h) - A similar pattern of polarized morphology was also induced by M4 TCM derived fractions containing ~ 12 kDa sized proteins (Fig. S5). Importantly, both GRO chemokines [69] and monomeric TGF-β [70] have a molecular weight of 12 kDa.

**Figure 7.**
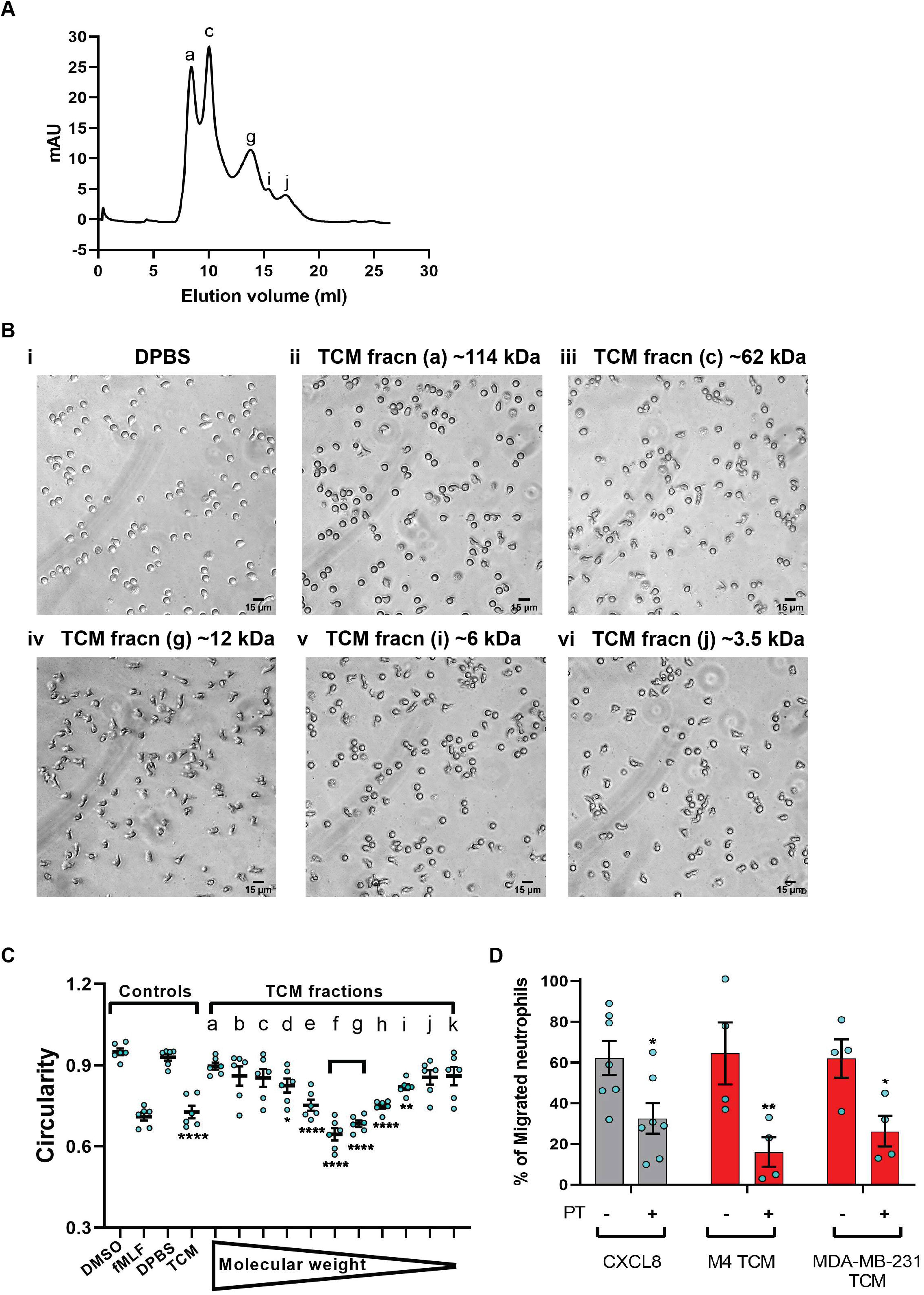
MDA-MB-231-derived active factors are ~12 kDa. **A**. Representative size exclusion chromatography elution profile of TCM from MDA-MB-231 cells. **B**. Representative bright field images of neutrophils exposed to vehicle control DPBS (Bi) or TCM peak fractions (fracn) (Bii-vi) as labeled in (A) and corresponding to different molecular weights. **C**. Graph depicting the circularity of neutrophils stimulated with controls (DMSO, fMLF, DPBS, TCM) or equal volume of TCM fractions with gradually decreasing molecular weight. Data show mean ± SEM from N=3 individual donors with two dots for each donor representing mean circularity of all neutrophils in two randomly selected imaging fields. **** P<0.0001, **P≤0.01, *P≤0.05 when compared with DPBS (one-way ANOVA with Dunnett’s multiple comparisons test). **D**. Graph depicting the percentage of neutrophils that migrated into the bottom chamber of the transwells containing equal volume of TCM or 20 nM CXCL8 in the presence or absence of PT. Bars show mean ± SEM from N=4-7 individual donors represented by each dot. **P≤0.01, *P≤0.05 when compared with corresponding PT-values (two-way ANOVA with Sidak multiple comparisons test).

Chemokines activate G protein coupled receptors (GPCR) by signaling through Gi heterotrimeric G proteins, which are sensitive to pertussis toxin (PT) inhibition [71]. To determine the role of chemokines in TCM induced neutrophil migration, we therefore, targeted chemokine specific GPCR Gi signaling by pretreating neutrophils with PT and measured the ability of cells to migrate towards TCM or CXCL8 as a positive control. Using transwell assays, we found that significantly less neutrophils migrated towards TCM derived from either M4 or MDA-MB-231 cells in the presence of PT (Fig. 7D), indicating a role for chemokines in TCM-induced neutrophil recruitment. PT treatment was not toxic, as it did not alter the migration of neutrophils towards the growth factor GM-CSF (Fig. S4). Taken together, these data suggest that chemokines actively contribute to TCM induced neutrophil migration.

### The neutrophil recruiting activity of the TCM depends on the combined action of GRO chemokines and TGF-β

In order to specifically determine the identity of the mediators, we studied TCM induced neutrophil migration in the presence of different inhibitors targeting either the GRO receptor CXCR2 (AZD 5069 [72]) or the TGF-β1 receptor ALK5 (SB431542 [73]), or both. The presence of both inhibitors did not alter neutrophil migration towards fMLF, showing that the inhibitors are receptor specific (Fig. 8A). We found that neutrophil migration induced by M4 TCM remained intact when cells were pretreated with either inhibitor alone (Fig. 8A). In contrast, we noted a significant decrease (~38%) in the percentage of migrating neutrophils towards M4 TCM when cells were pretreated with both inhibitors, indicating that CXCR2 ligands and TGF-β mediate the effects of the TCM (Fig. 8A). Similarly, we found a significant inhibition of M4 TCM induced neutrophil polarization as reflected by the increase in circularity in the presence of both the inhibitors compared to TCM alone (Fig. 8B). However, neutrophil migration induced by MDA-MB-231 derived TCM was not significantly decreased by the inhibitors (Fig. 8A).

**Figure 8.**
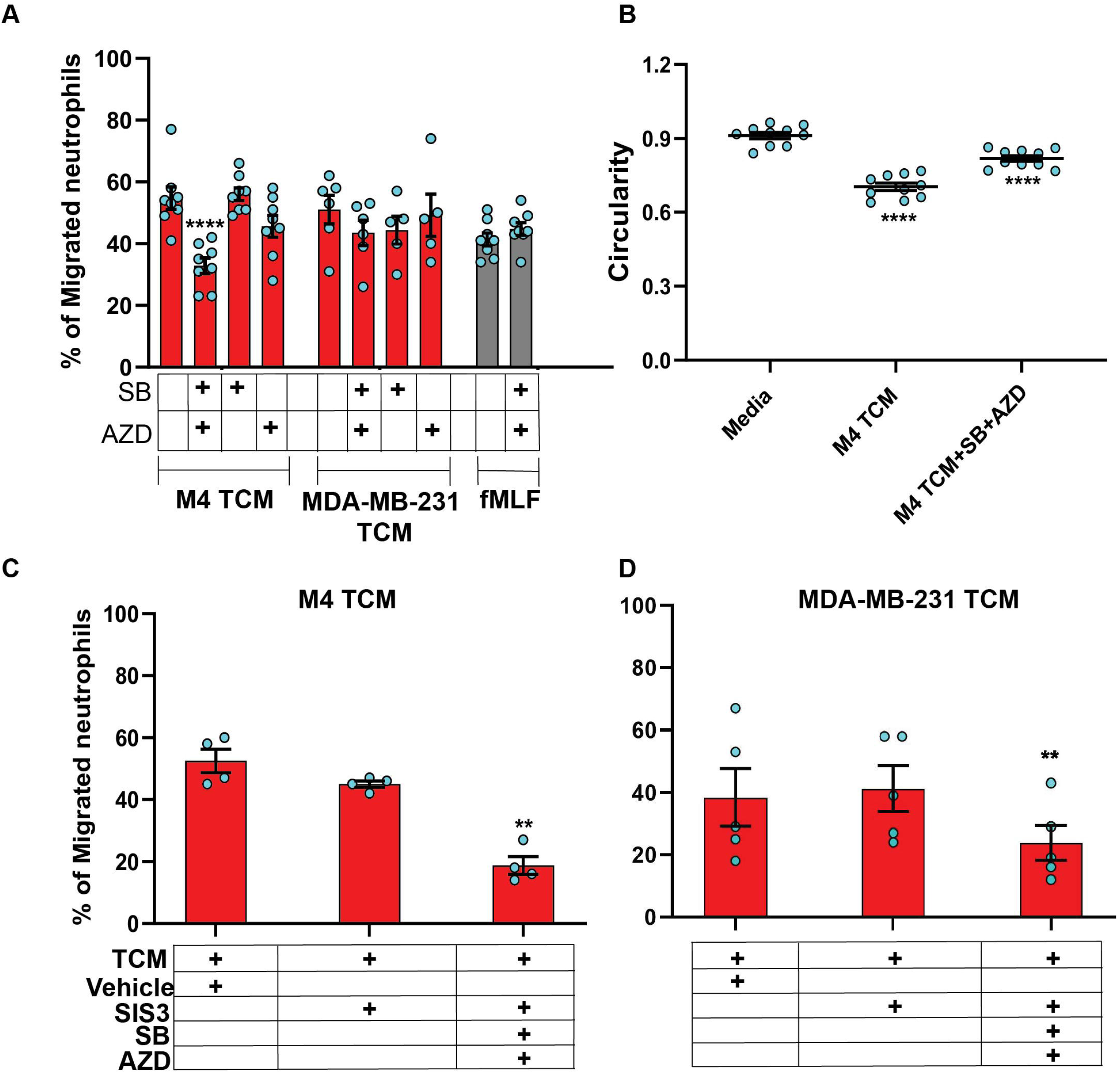
The neutrophil recruiting activity of the TCM depends on the combined action of GRO chemokines and TGF-β. **A**. Graph depicting the percentage of neutrophils either untreated or pretreated with 1 μM SB, 1 μM AZD, or a combination of 1 μM of SB and 1 μM AZD that migrated into the bottom chamber of the transwells containing equal volume of M4 TCM, MDA-MB-231 TCM, or 100 nM fMLF. Bars show mean ± SEM from N=5-8 donors with each dot representing individual donor. **B**. Graph depicting the circularity of neutrophils exposed to M4 TCM in the presence or absence of a combination of SB and AZD, or with media control. Bars show mean ± SEM from N=3 donors with 3-4 dots for each donor representing mean circularity of all neutrophils from at least 3 randomly selected imaging fields. **C&D**. Graphs depicting the percentage of neutrophils migrated into the bottom chamber of the transwells containing equal volume of M4 TCM (C) or MDA-MB-231 TCM (D) in the presence of 3 μM SIS3 alone or a combination of 1 μM each of SB, AZD and 3 μM SIS3, or vehicle control. Bars show mean ± SEM from N=4 (C) or 5 (D) donors. **** P<0.0001 when compared with untreated (A), media control (B) (one-way ANOVA with Dunnett’s multiple comparisons test). ** P≤0.01 when compared with vehicle control (C-D) (paired sample t-test).

To further elucidate the role of TGF-β signaling in mediating the effect of TCM on neutrophils, we targeted SMAD3, a downstream signaling component of TGF-β signal transduction pathway [74]. Neutrophils pretreated with SIS3 (an inhibitor that specifically inhibits SMAD3 phosphorylation [75]) showed intact migration in response to M4 TCM (Fig. 8C). However, when cells were pretreated with inhibitors targeting CXCR2, TGF-β receptor, and SMAD3 phosphorylation, we observed a remarkably stronger inhibition (~60%) of neutrophil migration towards M4 TCM (Fig. 8C) than the inhibition (~38%) measured with CXCR2 and TGF-β receptor inhibitors (Fig. 8A). Importantly, we also found a significant inhibition (~40%) of neutrophil migration towards MDA-MB-231 TCM in the presence of all three inhibitors, compared to vehicle control (Fig. 8D). Together, these results indicate that a concerted action of secreted TGF-β and GRO specific chemokines plays a key role in TCM induced neutrophil recruitment.

## Discussion

The purpose of this study was to determine whether neutrophil recruitment in breast tumors is dictated by the malignant potential of breast cancer cells. A number of retrospective studies have reported a unique association of neutrophils with TNBC compared to HR+ subtypes [16, 76, 77]. Using a panel of breast cancer cell lines representing highly aggressive TNBC or poorly aggressive ER+ breast cancer subtypes, we studied tumor driven neutrophil migration in three distinct migration assay systems. We showed a direct causal relationship between the metastatic potential of human breast cancer cells and their ability to recruit neutrophils, and conclude that TNBC subtypes induce robust neutrophil migration by secreting soluble chemical cues. More importantly, we profiled tumor secreted chemical cues with a number of biochemical, pharmacological and immunological approaches and conclude that a unique combination of GRO chemokines and TGF-β are responsible for driving neutrophil recruitment to breast tumor.

The primary function of GRO is to guide neutrophil navigate to infection or tissue injury sites. The GRO member CXCL1 has also been implicated to recruit neutrophils in tumor mouse models [52, 78]. We measured massive amounts of GRO family chemokines in TCM derived from TNBC cells compared to ER+ breast cancer cells. Yet, surprisingly, we found that TCM activity remains intact after targeting the GRO receptor CXCR2. While GRO chemokines are potent agonists of CXCR2, they can also bind and activate CXCR1, although they have lower affinity towards CXCR1 [79]. Furthermore, the rate of ligand induced CXCR1 internalization from the cell surface is relatively slower than that for CXCR2 [80]. Therefore, a potential role of GRO-CXCR1 signaling in mediating the neutrophil recruitment activity of TCM derived from TNBC cell lines cannot be excluded.

Factors/s other than chemokines may also play a key role to mediate neutrophil migration. In addition to GRO chemokines, we detected considerable amount of TGF-β1 cytokine in the TCM from both M4 and TNBC cell lines. The elevated presence of TGF-β in the tumor niche has been shown to facilitate tumor progression and metastasis in different cancer types [81, 82]. However, TGF-β lacks direct chemoattracting ability for neutrophils [83], which we also verified using recombinant TGF-β in a transwell setup (data not shown). Interestingly, TGF-β was recently shown to promote tumor-permissive phenotypes in TANs [84]. The same study also reported that TGF-β inhibition leads to massive infiltration of cytotoxic neutrophils in tumor-bearing mice. When we inhibited the TGF-β receptor TβRI/type I receptor ALK5, neutrophil migration induced by TCM remained intact. However, targeting both TβRI and CXCR2 led to a significant defect in neutrophil recruitment by M4 TCM, but not MDA-MB-231 TCM. Earlier studies identified a novel role of tumor-derived TGF-β in modulating chemokine receptor expression on immune cells [85]. Notably, tumor secreted factors were recently shown to increase expression of TβRI on neutrophils [86]. In line with these previous findings, we envision that TGF-β indirectly promotes TCM induced neutrophil migration through amplification of chemokine signaling. A key mediator in the canonical TGF-β signaling pathway that is critical for TGF-β1-triggered transcriptional regulation is Smad3 [74]. We found that addition of the Smad3 inhibitor disrupted the activity of both M4 and MDA-MB-231 TCM, supporting the notion that TGF-β signaling indirectly promotes neutrophil migration induced by chemokines isolated from TNBC cell lines. We envision that the different amounts of GRO chemokines and TGF-β in M4 and MDA-MB-231 TCM is responsible for the distinct sensitivity between the two TCM. It remains to be determined how TGF-β and chemokine signaling pathways regulate neutrophil migration.

While the transwell migration assay is ideal to quantify chemoattracting ability of multiple conditions simultaneously, it lacks the scope to directly monitor migrating cells and assess directionality of cell movement - a true indicator of chemotaxis. The neutrophil recruiting ability of tumor secreted factors collected from lung cancer cell lines has been reported using the underagarose assay [87], which allows to visualize and quantify cell migration [36]. However, we observed no neutrophil migration in response to M4 and MDA-MB-231-derived TCM using the underagarose setup (unpublished). The varying TCM content and diffusability of the secreted factors through the agarose could explain this discrepancy. We did find that the M4 TCM induced neutrophil chemotaxis in 3D collagen matrices that closely mimics interstitial space *in vivo*. By tracking migrating neutrophils, we showed that M4 TCM induced neutrophils to migrate with increased velocity and directionality, compared to media control. Using a co-culture system, we also showed that neutrophils migrate directionally toward M4 spheroids, while we measured no directional migration toward spheroids in a neutrophil-BT474 spheroid co-culture system. Cancer cells grown as 3D spheroids closely mimic the complexity of cell-cell, cell-matrix interactions, and gene expression profiles of the 3D tumor mass *in vivo* [88]. Others have shown neutrophil infiltration into lung tumor spheroids using endpoint imaging of tumor sections [89]. To our knowledge, our study is the first to demonstrate in real time that tumor spheroids actively recruit neutrophils. Whether neutrophils further amplify their chemotactic migration towards tumor spheroids through the secretion of secondary chemoattractants such as leukotriene B4 or other chemokines in response to tumor-derived factors remains to be addressed [24, 90]. Notably, we found that more neutrophils were present at the 20 hr endpoint in the vicinity of M4 spheroids, compared to BT474 spheroids. Neutrophils have very short life spans, however recent studies point to the role of pro-inflammatory soluble mediators in extending neutrophil survival [64, 91]. Indeed, many of the soluble factors including CXCL8, G-CSF, and GM-CSF, secreted abundantly from the M4 and MDA-MB-231metastatic breast cancer cells, are known to have pro-survival effect on neutrophils [64, 91]. Hence, we envisage that the pro-survival effect of M4 secreted factors also underlie the persistence of more neutrophils near M4 spheroids.

Much remains to be determined about the mechanisms underlying the secretion of active factors by highly malignant breast cancer cells. Others have shown a critical role of the epithelial-to-mesenchymal transition (EMT) program in promoting tumor metastasis [92, 93]. Through EMT epithelial cancer cells acquire mesenchymal features with enhanced invading properties that aid in cancer cell dissemination. Notably, myeloid cell recruitment was recently detected in EMT marker positive TNBC breast cancer specimen indicating possible role of EMT in tumor-driven neutrophil recruitment [94]. Moreover, EMT associated changes were connected to increased secretion of inflammatory soluble factors including CXCL8 from breast cancer cell lines [94]. Future studies will determine whether turning on EMT also enables breast cancer cells to secrete neutrophil recruiting GRO chemokines and TGF-β. Recently, the release of tumor-derived extracellular vesicles (EVs) have been suggested as a mechanism for the secretion of diverse soluble factors including chemokines and TGF-β [95, 96]. Whether similar mechanisms underlie the secretion of active factors by highly malignant breast cancer cells is the topic of future studies.

In conclusion, findings of this study complement previously established role of CXCL chemokines in mediating neutrophil recruitment to tumors, and expand our current understanding by directly linking malignant potential of breast cancer cells with their neutrophil recruiting ability. It is also the first study to unveil a novel role of tumor-derived TGF-β to cooperate with GRO chemokines and orchestrate neutrophil recruitment to TNBC tumors. Future studies will aim to understand how TCM impact the previously described pro-metastasis phenotypes of neutrophils [3, 9–14, 17, 97]. Knowledge from this study may help develop novel therapeutic strategies to target neutrophil recruitment to tumors and TAN function, which would restrict metastatic spread without perturbing protective functions of neutrophils against infection and injury.

## Abbreviations

TNBC: Triple negative breast cancer
ER: Estrogen receptor
HR: Hormone receptor
TAN: Tumor associated neutrophil
TCM: Tumor conditioned media
f-MLF: Formyl-methionyl-leucyl-phenylalanine
TGF-β: Transforming growth factor-β
GRO: Growth related oncogene
G-CSF: Granulocyte colony stimulating factor
GM-CSF: Granulocyte macrophage colony stimulating factor
EMT: Epithelial-to-mesenchymal transition
PT: Pertussis toxin

## Acknowledgements

We thank all the members of the Parent laboratory for their insights and suggestions. We are indebted to Dr. Sofia Merajver and Dr. James Rae for kindly providing cell lines. We thank Wu Zhi for thoughtful discussions and isolating TCM. We also thank Kalina Tsolova, Chiamaka Ukachukwu, and Kristen Loesel for assisting in the transwell, underagarose, and 3D migration assays, respectively. We acknowledge Dr. Michael Holinstat and Amanda Prieur from the Platelet Physiology and Pharmacology Core for providing blood draws for this study and thank Peilin Shen for neutrophil isolation and technical assistance. This work was supported by funding from the University of Michigan School of Medicine.

## Notes

### Competing Interest Statement

The authors have declared no competing interest.

