## Supplemental Information for "TRIPLE-NEGATIVE BREAST CANCER CELLS RECRUIT NEUTROPHILS BY SECRETING TGF-β AND CXCR2 LIGANDS"

**Supplemental Material**

### Supplementary Figure Legends

**Figure S1. The expression of CXCL1, CXCL2, CXCL5, and CXCL8 is elevated in breast cancer specimens with low ESR1 levels**

**A.** Representative heatmap from the Farmer study [40] with ESR1 and CXCL chemokines annotated with specific probeset ID. Datasets identified from the Oncomine platform were used to compare CXCL transcript expression between ER+ (blue) and ER- (red) breast cancer specimens sorted by ESR1 expression levels. The heatmap was generated by plotting log2 fold change of expression values relative to averaged ER+ for ESR1 and each of the CXCL chemokines. Blue and red indicate high and low expression, respectively.

**B.** Heatmap generated by plotting log2 fold change of averaged ER- relative to averaged ER+ for ESR1 and each of the CXCL chemokines calculated from the datasets of N=7 independent studies [40-46].

**C.** Summary of fold changes for CXCL1, CXCL2, and CXCL8 between ER- and ER+ with the adjusted P value from each of the 7 studies. A fold change of 1.5 or greater and an adjusted P-value of 0.05 or less was considered statistically significant.

**D.** Datasets from TCGA-BRCA were analyzed for ESR1 and CXCL chemokine expression. The log2 of the FPKM values were plotted for the chemokines and ordered by the FPKM of ESR1.

**Figure S2. TNBC cell lines have greater migration and invasion ability than ER+ breast cancer cells**

Graph depicting the migration (**A**) or invasion (**B**) potential of a panel of TNBC and ER+ cell lines. Results are depicted as mean ± SEM from N=3-5 independent experiments represented by each dot. **** P<0.0001, ** P≤0.01 when compared with values of MCF-7 cell line (one-way ANOVA with Dunnett’s multiple comparisons test).

**Figure S3. Active factors secreted from TNBC cell lines induce robust ERK phosphorylation**

Representative immunoblots depicting the levels of phospho-ERK and α-tubulin in response to fMLF, CXCL1, and TCM derived from M4, MDA-MB-231, and MCF-7 cells. Blot is representatives of four independent experiments.

**Figure S4. Treatment with PT specifically inhibits Gαi dependent signaling**

**A&B** Representative fluorescent endpoint images of neutrophils migrating underagarose towards a gradient 50 µg/ml GM-CSF in the absence or presence of pre-treatment with 500 ng/ml PT for 2 hrs.

**C&D** Representative fluorescent endpoint images of neutrophils migrating towards a gradient 100 nM fMLF in the absence or presence of pre-treatment with 500 ng/ml PT for 2 hrs. Images are representative of at least N=3 independent donors and acquired using an Axiovert fluorescence microscope from the side of the wells facing the chemoattractants (see Methods).

**Figure S5. M4-derived active factors are ~12 kDa**

**A**. Representative size exclusion chromatography elution profile of TCM from M4 cells.

**B**. Representative bright field images of neutrophils exposed to vehicle control DPBS (Bi) or TCM peak fractions (fracn) (Bii-vi) as labeled in (A) and corresponding to different molecular weights.

**C**. Graph depicting the circularity of neutrophils stimulated with controls (fMLF, DPBS, TCM) or TCM fractions with gradually decreasing molecular weight. Data show mean ± SD from N=1 donor with two dots representing mean circularity of all neutrophils from two randomly selected imaging fields.

**Supplementary Movie Legends**

**Movies S1&S2** Videos showing time-lapse imaging of labeled neutrophils migrating in 3D collagen matrices in response to M4 TCM (S1) or media (S2) placed on the right. Images were taken every 25 sec over 40 min. Individual cell tracks in the immediate vicinity of TCM-collagen boundary are shown.

**Movies S3&S4** Videos showing time-lapse imaging of labeled neutrophils migrating in 3D collagen matrices in response to 500 nM CXCL8 (S3) or buffer (S4) placed on the right. Images were taken every 25 sec over 50 min. Individual cell tracks in the vicinity of TCM-collagen boundary are shown.

**Movies S5-S8** Videos showing z-stacks of 2-photon images of M4 (Total slice no. 38; S5), MDA-MB-231 (Total slice no. 23; S6), BT549 (Total slice no. 31; S7), and BT474 (Total slice no. 147; S8) spheroids stained with a nuclear dye DRAQ5.

**Movies S9&S10** Videos showing time-lapse imaging of the neutrophil-M4 (S9) and -BT474 (S10) spheroid co-culture system. Live-imaging was initiated one hour after the addition of neutrophils to the spheroids. Images were taken every 3 min for 20 hrs. Individual neutrophil tracks are shown.
