## Supplementary figures and images for "TRIPLE-NEGATIVE BREAST CANCER CELLS RECRUIT NEUTROPHILS BY SECRETING TGF-β AND CXCR2 LIGANDS"

### Supplemental Figure 1

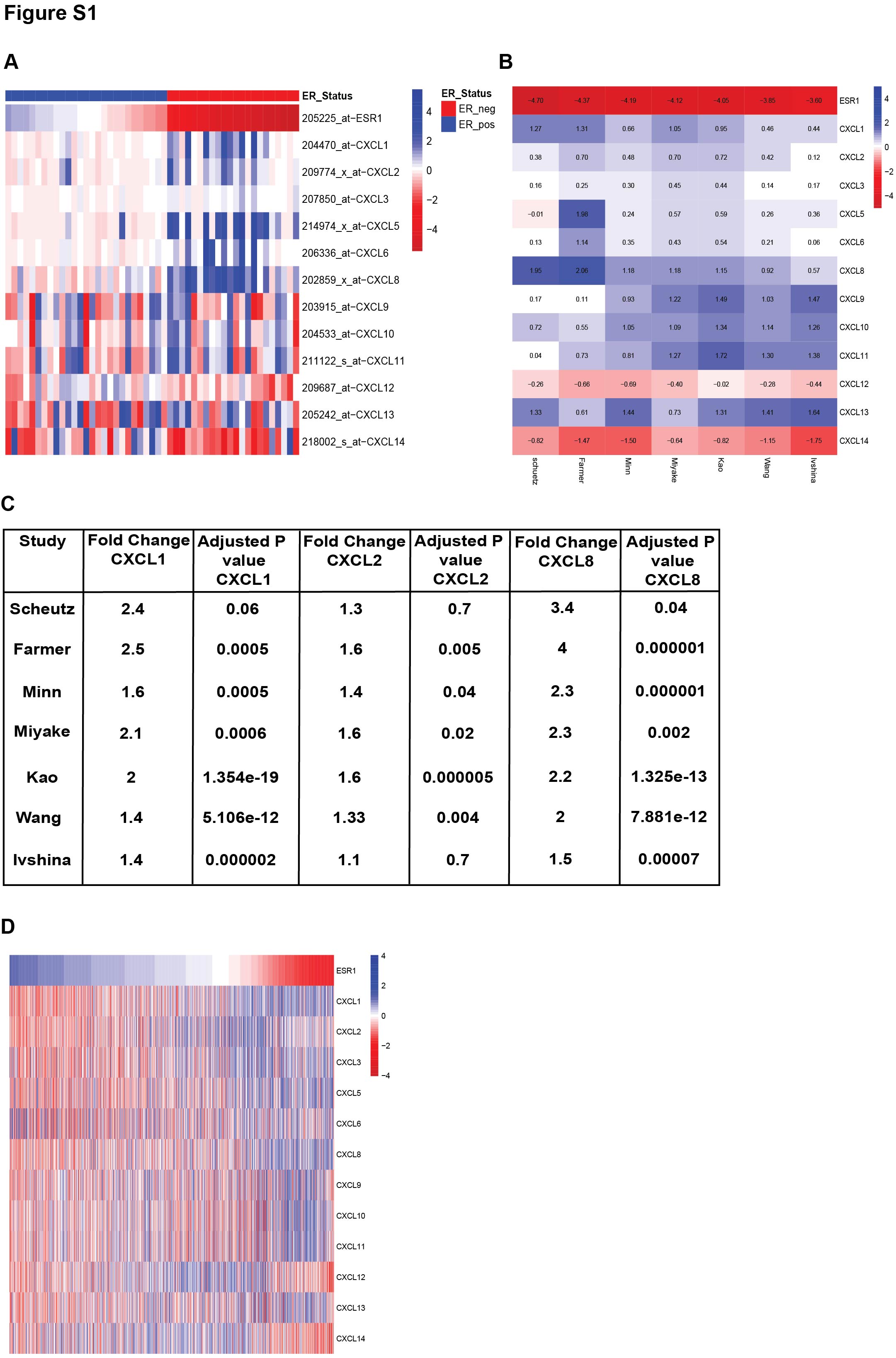

### Supplemental Figure 2

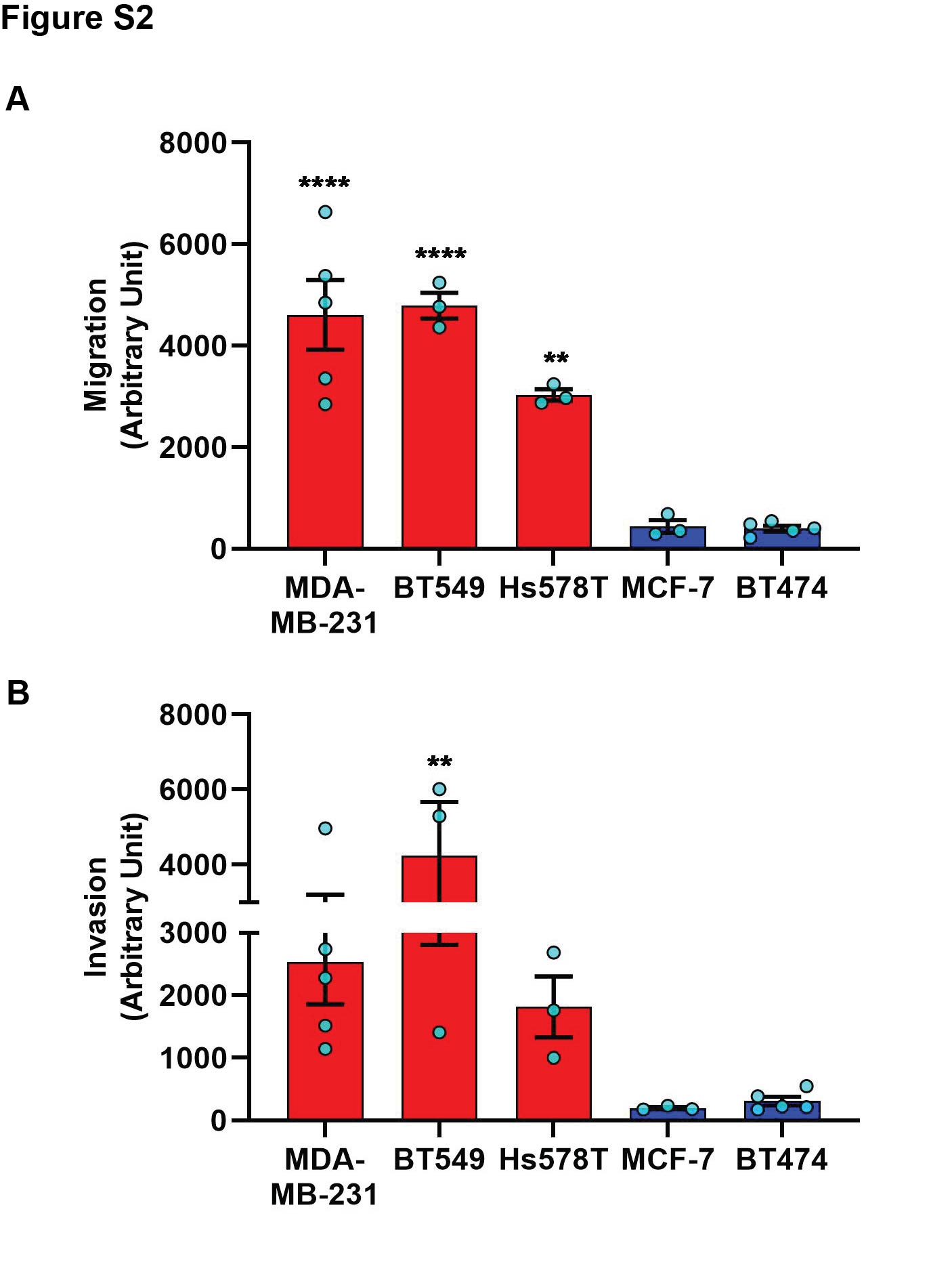

### Supplemental Figure 3

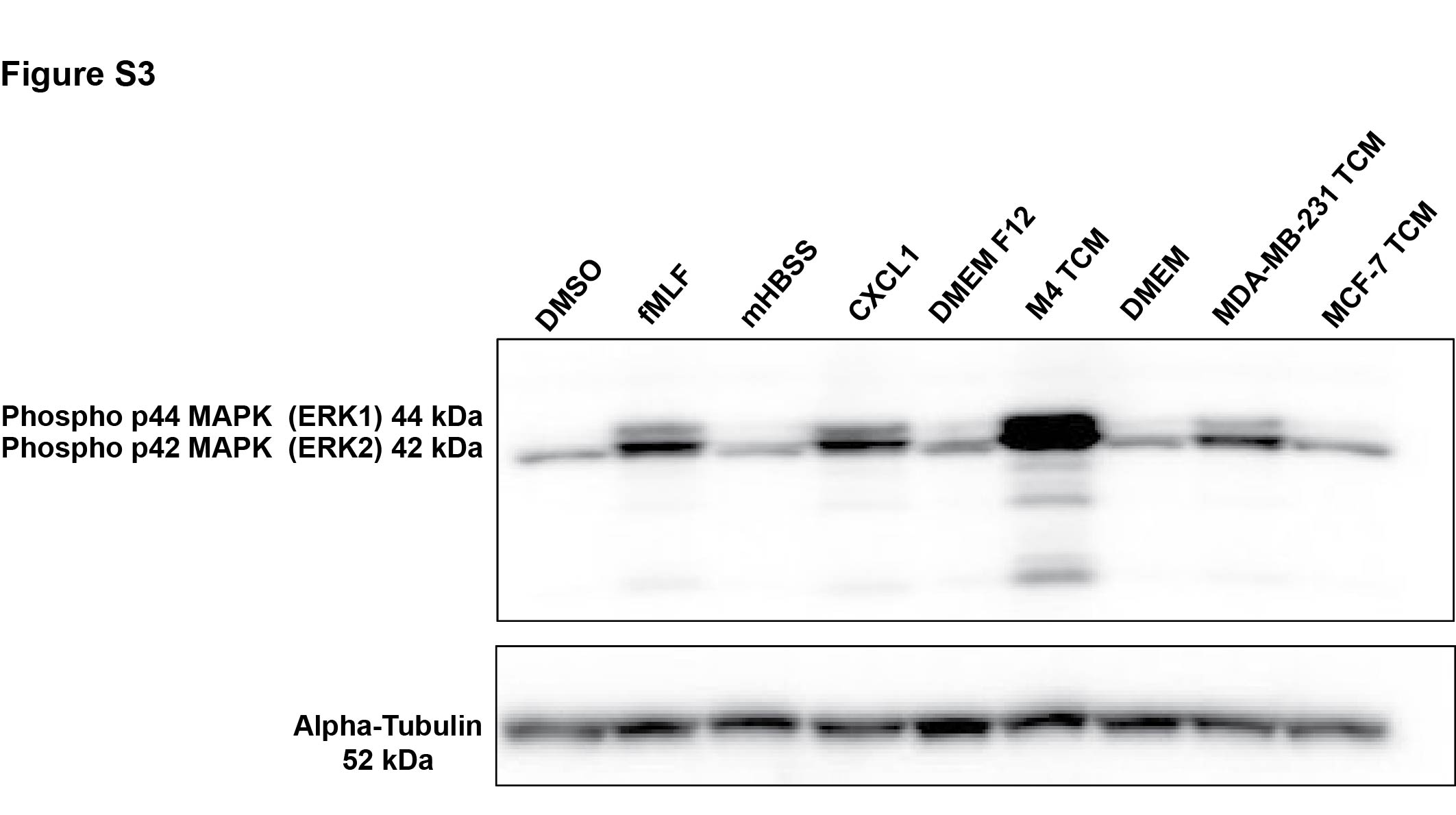

### Supplemental Figure 4

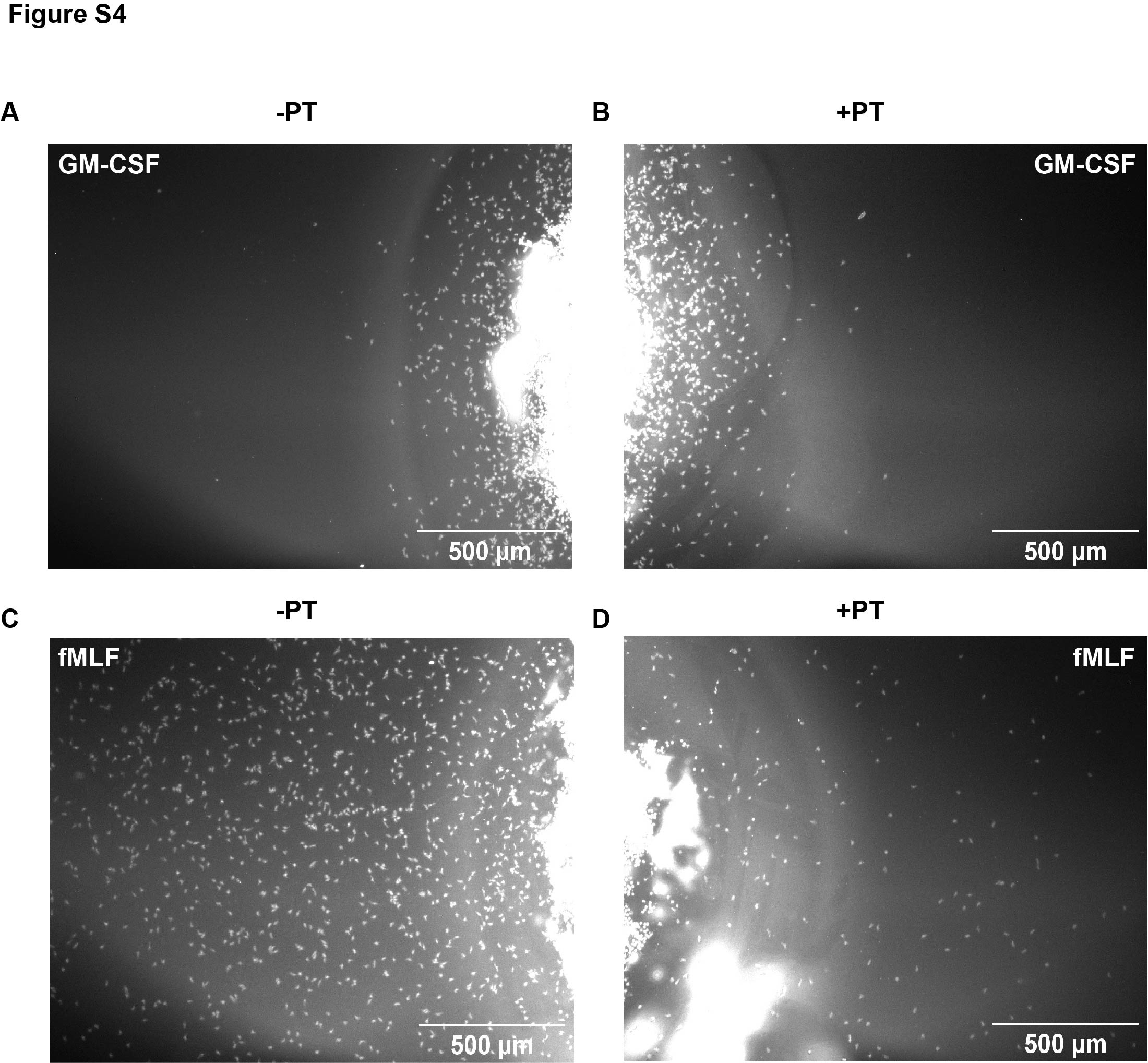

### Supplemental Figure 5

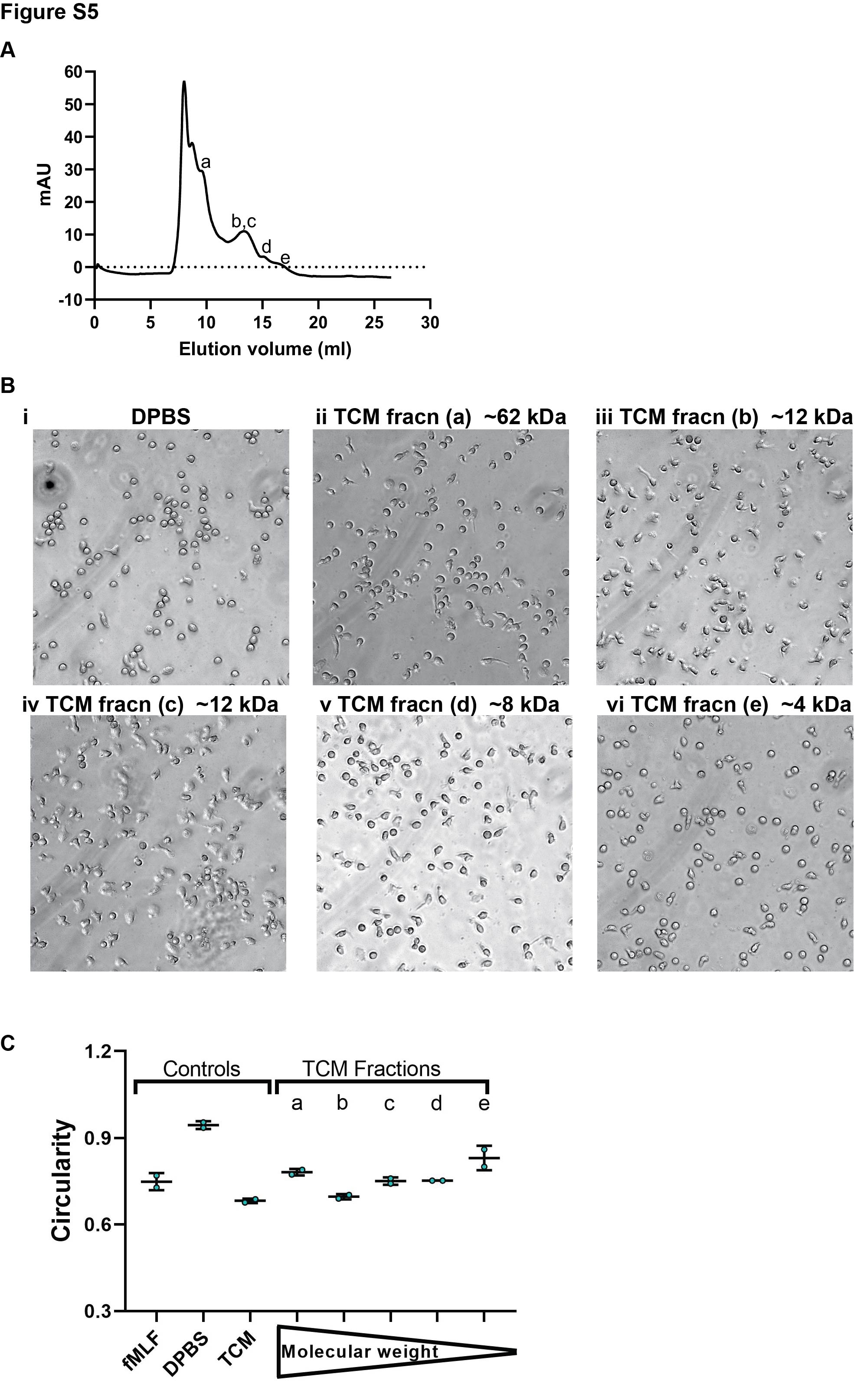
